# Equipositioning of Chromosomes in the Polyploid Archaeon *Haloferax volcanii* by HpaAB

**DOI:** 10.1101/2025.11.19.689047

**Authors:** Kartikeyan Premrajka, Ulrike Johnsen, Holger Preidel, Jascha Wienberg, Chiara Corleis, Marius Ortjohann, Giacomo Giacomelli, Marc Bramkamp

## Abstract

Proper genome segregation is essential for all life forms, but in archaea, this process is less characterized as compared to bacteria, or eukarya. Archaea show considerable variability in chromosome and plasmid copy number, and organization. Stable maintenance of chromosomes and low-copy plasmids in organisms requires an active segregation system. *Haloferax volcanii* DS2 is a polyploid archaeon that contains a fragmented genome composed of a main chromosome, three secondary chromosomes, pHV1, pHV3 and pHV4, as well as a plasmid pHV2. Due to the high copy number of all replicons, genomic segregation in *H. volcanii* is typically considered random. However, we identified four two-gene operons encoding a ParA-like homologue and an uncharacterized protein. Here we characterized one of these systems, termed here HpaA (ParA-homolog) and HpaB, that is encoded on the secondary chromosome pHV3. HpaA, showing conserved ATPase walker motifs, is a ParA protein with a bacterial origin acquired via horizontal gene transfer. Mutation of key residues in HpaA alters its localization, and HpaA is necessary for proper spatial distribution of HpaB. HpaB, an archaeal protein capable of forming foci in the cytosol, is shown to function as a centromere-binding protein, thereby acting as a functional ParB homologue. qPCR data confirm that HpaB is crucial for maintaining the plasmid-to-chromosome ratio, as deletion of *hpaB* reduces pHV3 copy number. Single particle tracking suggests that absence of HpaB leads to an altered population distribution of HpaA, based on protein dynamics. Deletion of the genes *hpaA and hpaB* results in severe growth defects when cells are grown on xylose as a sole carbon source, in line with the location of the genes for the xylose degradation pathway on chromosome pHV3. We propose that *H. volcanii* has an active chromosome segregation system, essential for the “two-dimensional” segregation and equipositioning of its replicons. This function is crucial for ensuring stable replicon maintenance across the distinct cell morphologies of *H. volcanii*, which transition between rod-shaped and disc-shaped forms at different stages of growth.

## Introduction

Proper segregation of the genome is a fundamental process across all domains of life, ensuring the accurate transmission of genetic material to progeny during cell division. While this process has been extensively characterized in both Eukarya ^1,2^ and Bacteria ^3-6^, our understanding of genome segregation in Archaea remains limited. In bacteria, one of the most comprehensively studied DNA segregation systems is the ParABS system. Originally identified in plasmid partitioning ^3,7-11^, it was later found to also function in chromosomal segregation across diverse bacterial species ^4,12,13^. The system comprises three core components: ParA, an ATPase; ParB, a site-specific DNA-binding protein; and *parS*, a centromere-like DNA sequence ^12,14^. ParB recognizes *parS* sites that are typically clustered around the origin of replication (*oriC*) through its CTP-binding and DNA-binding domains ^12,15-18^. ParB forms dimers with high affinity for *parS* ^4,12^ followed by CTP binding of the *parS*-bound ParB ^19,20^ which allows for clamping of ParB onto the DNA and liberates it from *parS* allowing spreading onto proximal DNA ^21^. Subsequent CTP hydrolysis triggers unclamping and release of ParB from DNA, allowing for recycling and maintaining a dynamic pool of ParB molecules around *oriC* ^22^. This CTPase activity is essential for forming the partitioning complex ^17,19,20^. Additionally, ParB has been shown to be capable of CTP-dependent liquid-liquid phase separation (LLPS) creating condensates that concentrate ParB at *parS* sites. ParA-ATP driven interactions allows for positioning or splitting these condensates, in turn supporting efficient segregation ^23-25^.

ParA, a Walker-type ATPase, binds non-specifically to the nucleoid DNA in its ATP-bound dimeric form ^26-29^. Interaction with *parS*-bound ParB triggers ATP hydrolysis, locally releasing ParA from the DNA and generating a dynamic ParA distribution ^30^. The resulting spatial asymmetry of ParA drives a diffusion-ratchet mechanism, whereby ParB-*parS* complexes move toward regions enriched in DNA-bound ParA ^30-34^, producing directional segregation and bipolar positioning of *parS* containing genetic elements. The ParABS system has been characterized in a variety of bacteria, including *Bacillus subtilis* ^35^, *Corynebacterium glutamicum* ^36^, *Caulobacter crescentus* ^30,37^ and *Myxococcus xanthus* ^13^.

Interestingly, the ParA family of ATPases also contain multiple proteins that are involved in cargo positioning or spatial regulation. For instance, gradient based mechanisms allow distribution of cargo within the cell, such as carboxysome positioning via McdA oscillation ^38^. On the other hand, division site positioning is regulated by MinD gradients in *E. coli* ^39^, while a bipolar gradient is formed by MipZ ^40^ in alpha proteobacteria like *Caulobacter*. Interestingly, although ParB stimulates ParA’s ATP hydrolysis in some systems, its interaction with MipZ instead promotes MipZ dimerization, generating a distinctly different gradient from that of ParA.^40^.

In contrast to bacteria, the mechanisms of DNA segregation in most archaeal lineages remain poorly understood. Some progress has been made in model crenarchaeal species such as *Sulfolobus solfataricus* and *Sulfolobus* strain NOB8H2. Two systems, SegAB ^41,42^ and AspA-ParBA ^43^, have been implicated in chromosome and plasmid segregation in these organisms, respectively. The AspA-ParBA system, encoded on the pNOB8 plasmid of *Sulfolobus NOB8H2*, features components with varying degrees of similarity to bacterial homologs. The ParB and ParA proteins share 42–58% and 33–37% sequence identity with their bacterial counterparts, respectively ^44^. However, AspA is unique to Archaea and contains a helix-turn-helix motif. AspA binds specifically to a 23 bp palindromic sequence upstream of the segregation cassette and spreads along adjacent DNA at high concentrations, forming a DNA-protein superhelix ^43^. ParB in this system functions as a bridging adaptor between AspA and ParA. It has a modular structure, with the N-terminal domain binding AspA, the C-terminal domain binding non-specifically to DNA, and a linker region responsible for ParA interaction. ParA, like its bacterial counterpart, binds DNA in an ATP-dependent manner. The *aspA-parB-parA* cassette is widespread among Crenarchaeota, appearing on both plasmids and chromosomes ^43^.

Archaea are a genetically diverse domain, and cell division mechanisms, genome ploidy and organization vary widely among its lineages ^45^. For example; Euryarchaea have been shown to be oligoploid ^46,47^, while Crenarchaea have one or two copies of their chromosomes in different stages of the cell cycle ^48,49^. One major challenge in characterizing archaeal DNA segregation stems from their varied ploidy. For instance, the halophilic archaeon *Haloferax volcanii* contains approximately 20 chromosomal copies during exponential growth and 12 in stationary phase ^50^, along with secondary chromosomes ^51^. High ploidy in Euryarchaea, including *H. volcanii*, coincides with the presence of histones ^52^, facilitating DNA compaction. In *H. volcanii* strain DS2, the genome comprises a 2.8 Mb main chromosome, three secondary chromosomes pHV4 (636 kb), pHV3 (438 kb), and pHV1 (85.1 kb) and the pHV2 plasmid (6.35 kb) ^53,54^. All secondary chromosomes contain a single origin of replication, while the main chromosome contains three origins ^46,53,55^. However, in the laboratory strain H26 used in this study, pHV2 has been cured ^56,57^, and pHV4 is stably integrated into the chromosome due to a fusion event ^46^. It is generally believed that H. volcanii relies on random distribution rather than an active segregation mechanism for its chromosome, due to its high copy number and lack of identifiable centromere-like structures ^58^. However, the segregation of secondary chromosomes and plasmids, which are maintained in equimolar ratios with the main chromosome (1:1 to 1:1.2), remains poorly understood.

In this context, we identified several ParA-like genes throughout the *H. volcanii* genome, all organized in putative operons with adjacent uncharacterized genes. One such operon, located on the secondary chromosome pHV3, contains genes HVO_B0018 (Haloarchaeal Partitioning protein A - HpaA) and HVO_B0017 (Haloarchaeal Partitioning protein B - HpaB). While HpaA contains the conserved “deviant Walker motif”, which is a hallmark of canonical ParA/MinD like ATPases ^59^, and the operon organization strongly resembles those of previously described *parABS* systems, HpaB shows no sequence similarity to conventional ParB proteins.

Given the domain structure and gene organization, along with the established role of ParA-family proteins in cargo positioning ^60^, we hypothesized that HpaA-HpaB constitutes a novel chromosome segregation system in *H. volcanii* in which a ParA-like protein relies on a novel class of chromosomal ParB functional homologues. We present evidence that the HpaAB system operates as an equidistant positioning mechanism, maintaining uniform spacing between chromosomes. This is mechanistically slightly different to a classical segregation system that allows directional transport of DNA cargo. Moreover, equipositioning allows effective segregation in both morphologies of *H. volcanii* – rods in early growth phases, and plates in late growth phases.

The unique physiology of Archaea and the presence of hybrid segregation systems such as AspA-ParBA and the here described HpaAB, offers a window into the evolutionary origins of cellular organization, particularly with respect to Horizontal Gene Transfer (HGT). HGT allows for gene exchange between prokaryotes ^61^. Haloarchaea, including *H. volcanii*, have acquired a substantial portion of their genome from bacteria ^62,63^. Understanding how secondary chromosomes and plasmids are inherited and maintained in such highly polyploid and pleiomorphic organisms is therefore critical. It will not only shed light on genomic stability in complex prokaryotes but also on the evolution of ATPase-based cargo positioning systems across the domains of life.

## Material & Methods

### Growth Conditions and Media

All strains are based on the *H. volcanii* H26 serving as the wild type (Supplementary Material Table S1). All strains were grown at 42° C under aerobic conditions in synthetic medium ^64^ supplemented with either casamino acids (1 % w/v), D-glucose (20 mM) or D-xylose (20 mM). The wild type and deletion strains that did not carry a plasmid were grown in the presence of uracil (50 μg/ml). Strains transformed with either the empty vector pTA963 ^65,66^ or derived plasmids were grown in medium without the addition of uracil to select for plasmid carrying cells. Cells were always pre-inoculated directly from glycerol stocks into media containing 1% casamino acids for 24-48 h and then transferred to media containing the desired carbon source.

Growth of the wild type and of the *ΔhpaA* and *ΔhpaB* deletion strains was compared on D-xylose. For complementation of the uracil auxotrophy, these strains carried the empty vector pTA963. For in trans complementation, the deletion mutants were transformed with plasmids derived from pTA963, encoding the corresponding genes (*hpaA* or *hpaB)* under the control of the tryptophan inducible promotor p.tnaA. Precultures of the strains were grown on 1% casamino acids supplemented with 0.1 mM tryptophan. Growth experiments were performed in 100 ml flasks containing 20 ml synthetic medium supplemented with 20 mM xylose and 0.1 mM tryptophan by measuring the optical density at 600 nm (OD_600_). Growth was started at an OD_600_ of 0.05 and followed for approximately 80 h.

### Generation of deletion mutants

All strains, plasmids and oligonucleotides are listed in supplementary material tables S1-3. Gene deletion mutants of *hpaA*, and *hpaB* were constructed using the pop-in/pop-out strategy ^57^. The 500 base pairs each upstream and downstream of the genes, excluding the genes, were fused and cloned into the suicide vector pTA131 using Gibson’s assembly ^67^. *H. volcanii* H26 cells were transformed with the plasmids after multiplication in E. coli (XL1 Blue MRF). *H. volcanii* H26 was transformed using polyethylene glycol 600 ^68^. Pop-in cells, which contain the plasmid integrated into the host genome via homologous recombination were selected via growth of colonies on agar plates without uracil, containing 1% casamino acids. Correct clones were verified for presence of plasmid via colony PCR, and passaged 5 times in synthetic medium to allow allelic integration into all copies of the genome. Pop-out clones were subsequently generated by plating on agar plates containing both uracil and 5-fluoroorotic acid (5-FOA) (50µg/ml). 5-FOA acted as the selective agent, as it is toxic for cells containing the *pyrE2* gene. Cells that had undergone a second homologous recombination event (pop-out) were able to grow, leading to either wild type or deletion variants which were screened and selected via colony PCR.

### Generation of strains expressing fusion proteins

Plasmids were constructed via restriction enzymes-based insertion of genes of interest, or via Gibsons cloning ^67^. Polymerase chain reactions were carried out using Q5 or Phusion® HF polymerase (New England Biolabs) according to the manufacturer’s instructions. Purified genomic or plasmid DNA served as the template. Primers with overlaps were designed via NEBuilder (NEB) for fusion proteins and Gibson assembly constructs. Point mutations, and indels were introduced using mutated primer overlaps via Gibson assembly, or via Q5 site-directed mutagenesis (NEB). Primer design for the latter was done via the NEBaseChanger tool (NEB). Plasmid pTA963^65^ was used as a basis to construct controlled expression of fluorescently tagged HpaA or HpaB variants. For plasmids that were constructed via Gibsons assembly (Gibson et al, 2009), the proteins of interest where amplified from the genome and fused with a fluorophore of interest via a GGGGSGGGGS linker. The fusion protein was controlled via the tryptophan inducible promotor p.tnaA. Induction was performed with 1 mM tryptophan for 1 hour when necessary.

### Fluorescence Microscopy

For still images, fluorescence microscopy was performed with exponentially grown cells mounted on agarose coated slides (1 % agarose in growth medium). Cultures were grown in synthetic media to an OD_600_ of 1.0 to obtain plate morphologies, and spotted on agarose pads and allowed to dry in room temperature followed by covering with a glass coverslip. Images were acquired on an Axio-Observer 7 fluorescence microscope (Carl Zeiss) equipped with a Colibri 7 LED light source (Zeiss) with an EC Plan Neofluar 100 x/1.3 oil Ph3 objective, a 2.5 x optovar and a Hamamatsu ORCA-R2 camera. msfGFP was visualized with a filter set 38 HE (BP470/40 FT 495 BP 525/50) and excited at 488 nm (100 % Colibri LED intensity) for 400 ms, mScarlet-I was visualized with a filter set 43 HE (BP550/25 FT570 BP605/70) and excited for 500 ms, while mTurqoise2 was visualized with a filter set 47 HE (BP 436/25 FT 455 HE BP 480/40) and excited for 100 ms. The microscope was equipped with an environmental chamber set to 42 °C. Digital images were acquired with Zen Blue (Zeiss) and analysed and edited using FIJI (version 2.17)^69^.

Time-lapse experiments were performed on exponentially grown cells after dilution to an OD600 of 0.05 in synthetic media and loaded in a microfluidic chamber (B04A CellASIC®, Onix); the environmental chamber was heated to 42° C and 0.75 psi were applied for nutrient supply throughout the experiment. Images were taken in 3-6 min intervals. Imaging of live cells was performed with the same setting used for imaging for cells mounted on agarose.

Computational analysis of agarose mounted cells was performed using the MicrobeJ plugin ^70^ from the image analysis platform FIJI (Version 2.17). MicrobeJ algorithm was used for automatic detection of cell (using restricted cell length and area to recognize real cells) and foci for extraction of corresponding regions of interest from all image channels. The regions of interest (cell contours, and foci) were converted into a map of X and Y coordinates. Data were further evaluated via these X and Y coordinates using the software R ^71^ and Rstudio (Version 4.5) ^72^.

### Super Resolution Microscopy and Single Particle Tracking

Exponentially growing *H. volcanii* WT and *ΔhpaB* cells expressing HpaA-HaloTag ectopically were stained with 50 nM HaloTag TMR Ligand (Promega) for 30 min at 42°C at 600 rpm on a standard 2 ml heat block (Eppendorf). Cells were harvested (6000 rpm, 2 min, RT) and washed three times in filtered synthetic media.

Slides were cleaned with 1 M KOH overnight, rinsed with deionized H_2_O and dried with pressurized air. Agar pads where prepared with the aid of gene frames (Thermo Fisher) and 1% (w/v) low melting agarose (agarose, low gelling temperature, Sigma-Aldrich) in sterile filtered synthetic media (0.2 μm pores).

Single molecule localization microscopy (SMLM) was performed with an Elyra 7 (Zeiss) inverted microscope equipped with pco.edge sCMOS 4.2 CL HS cameras (PCO AG), connected through a DuoLink (Zeiss), only one of which was used in this study. Cells were observed through an alpha Plan-Apochromat 63×/1.46 Oil Korr M27 Var2 objective in combination with an Optovar 1 × (Zeiss) magnification changer, yielding a pixel size of 97 nm. During image acquisition, the focus was maintained with the help of a Definite Focus.2 system (Zeiss). Fluorescence was excited with a 561 nm (100 mW) laser, and signals were observed through a multiple beam splitter (405/488/561/641 nm) and laser block filters (405/488/561/641 nm) followed by a Duolink SR DUO (Zeiss) filter module (secondary beam splitter: LP 560, emission filters: LP570). For each time lapse series, 10 000 frames were taken with 20 ms exposure time (∼24 ms with transfer time included) and 50% 561 nm intensity laser in TIRF mode (62° angle).

For single-particle tracking, spots were identified with the LoG Detector of TrackMate v6.0.1 ^73^, implemented in Fiji 1.53, an estimated diameter of 0.5 μm, and median filter and sub-pixel localization activated. The signal-to-noise threshold for the identification of the spots was set at 7. To limit the detection of ambiguous signal, frames belonging to the TMR bleaching phase (first 1000 frames) were removed from the time lapses prior to the identification of spots. Spots were merged into tracks via the Simple LAP Tracker of TrackMate, with a maximum linking distance of 500 nm, two frame gaps allowed, and a gap closing max distance of 0 nm. Only tracks with a minimum length of 5 frames were used for further analysis.

To identify differences in protein mobility and/or behaviour, the resulting tracks were subjected to mean-squared-displacement (MSD), square displacement (SQD) analysis, as described previously ^74^. Both the analytical approaches relied on SMTracker 2.0 ^75^.

The average MSD was calculated for four separate time points per strain (exposure of 20 ms—τ = 24, 48, 72 and 96 ms), followed by fitting of the data to a linear equation. The last time point of each track was excluded to avoid track-ending-related artefacts. The cumulative probability distribution of the square displacements (SQD) was used to estimate the diffusion constants and relative fractions of up to three diffusive states (‘fast’, ‘slow’, and ‘static’) ^76^.

### qPCR

To isolate genomic DNA, *H. volcanii* cells were grown to an OD_600_ of approximately 1.0. The genomic DNA was then extracted from the cells using the Wizard® Genomic DNA purification kit (Promega) according to the manufacturers’ manual with the following changes. After the centrifugation with the protein precipitation buffer to separate the cell debris, the DNA was pipetted twice with a 1 ml pipette tip to get a homogenous solution, then the supernatant was incubated in isopropanol overnight at -20° C. The concentration and purity of the isolated genomic DNA was verified using a nanophotometer wherein the A260/A280 ratio was around 1.8, and used for qPCR or stored at -20° C.

DNA amplification was performed using 2x qPCR Master mix SYBR blue (GeneON, Y220) according to the manufacturer’s manual. Reactions for plasmid and chromosome contained 200 nM of the respective oligonucleotides and 5 ng genomic DNA (total reaction volume of 25 µl). Samples were measured in technical triplicates via an AriaMx and *C*_t_ values were determined using the Agilent Aria Software 1.8v. Oligonucleotides efficiencies were estimated by calibration dilution curves and slope calculation; data were analysed by 2^−Δ^*C*^T^ method accounting for dilution factors and sample volumes used for DNA purification.

### Structure prediction

Alphafold 3 ^77^ was used to predict all structures of HpaA and HpaB variants in Figures 1, 8, Supplementary figure 1, and Supplementary Figure 4 The same seed was specified for all mutants of HpaB.

**Figure 1.**
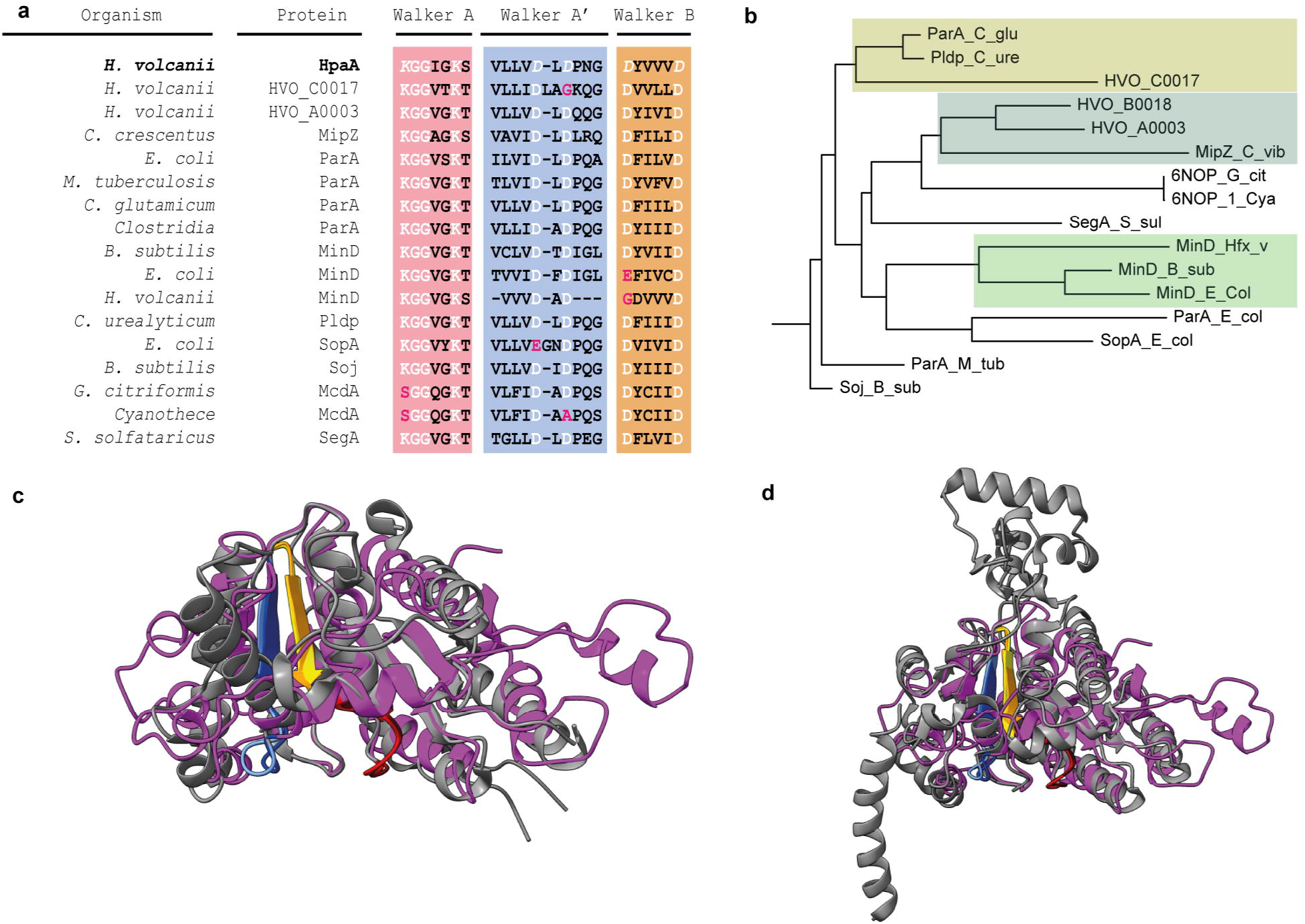
HpaA is a ParA homologue. (A) Alignment of the amino acid sequences of the Walker A, A’, and B motifs conserved among known bacterial and archaeal ParAs. The signature lysine, invariant glycines, catalytic lysine in the Walker A motif, and the catalytic aspartate in the Walker A ‘motif are conserved (highlighted in white). (B) Phylogenetic tree using the whole gene sequence of HpaA with other *Haloferax HpaAs* and known bacterial ParAs; made using neighbour-joining method. Colours indicate closer related proteins by branch length. The predicted structure of the AlphaFold model of the HpaA dimer in magenta when compared with known structures of MipZ (2XJ4) in (C) and ParA in (D) in gray shows an overlap of the walker motifs (Red/Dark Red - Walker A, Blue/Turquoise - Walker A’, Orange/Yellow - Walker B).

### Statistical analysis

All statistical analyses for cell morphologies, qPCR pHV3 to main chromosome ratios, nearest neighbour distance distributions, and linear regressions were performed in R (version 4.5) ^71^ and Studio (version 2025.09.2+418) ^72^. The codes are all available as supplementary material, with the appropriate packages cited. Morphological parameters (cell length, width), nearest neighbour distances of foci as x and y coordinates on a cartesian plane, and cell profiles were extracted for each individual cell from FIJI using the plugin MicrobeJ ^70^. To test for differences among strains (wild type, *ΔhpaA*, and *ΔhpaB*), Kruskal–Wallis test was used for each parameter, as the data were not normally distributed (Shapiro–Wilk test, p < 0.05). When the Kruskal–Wallis test indicated a significant difference (p < 0.05), Dunn’s post-hoc tests with Benjamini–Hochberg false discovery rate (FDR) correction were performed to compare each mutant strain against the wild type (WT). Adjusted p-values were used to determine statistical significance and summarized as follows: p < 0.05 (*), p < 0.01 (**), and p < 0.001 (***). Effect sizes were determined via Vargha and Delaney’s A. All tests were two-tailed unless otherwise noted. Statistical values are summarized in supplemental material table S4.

## Results

### The hpaAB operon comprises of an archaeal ParA homologue and a unique uncharacterized partner

*H. volcanii* H26 contains 4 ORFs encoding a ParA-like protein on its secondary chromosomes and plasmids; two on pHV4 and one each on pHV3 and pHV1 (Table 1). All the ORFs are each arranged in an operon where the genes of annotated ParA-family proteins are immediately upstream of genes of unknown function, with an overlap of 4-8 nucleotides. Moreover 3 out of the 4 operons are located in the vicinity of the origins of each secondary chromosome, which has been shown to be the case for *parAB* operons in the ParABS systems ^78,79^ which is likely due to the vicinity of the *parS* site to the origin ^6^. Only pHV4, which has integrated into the main chromosome in the lab strain H26 has two such operons with the second (HVO_A0458 and HVO_A0459) being distant from the *ori* of pHV4.

**Table 1.**
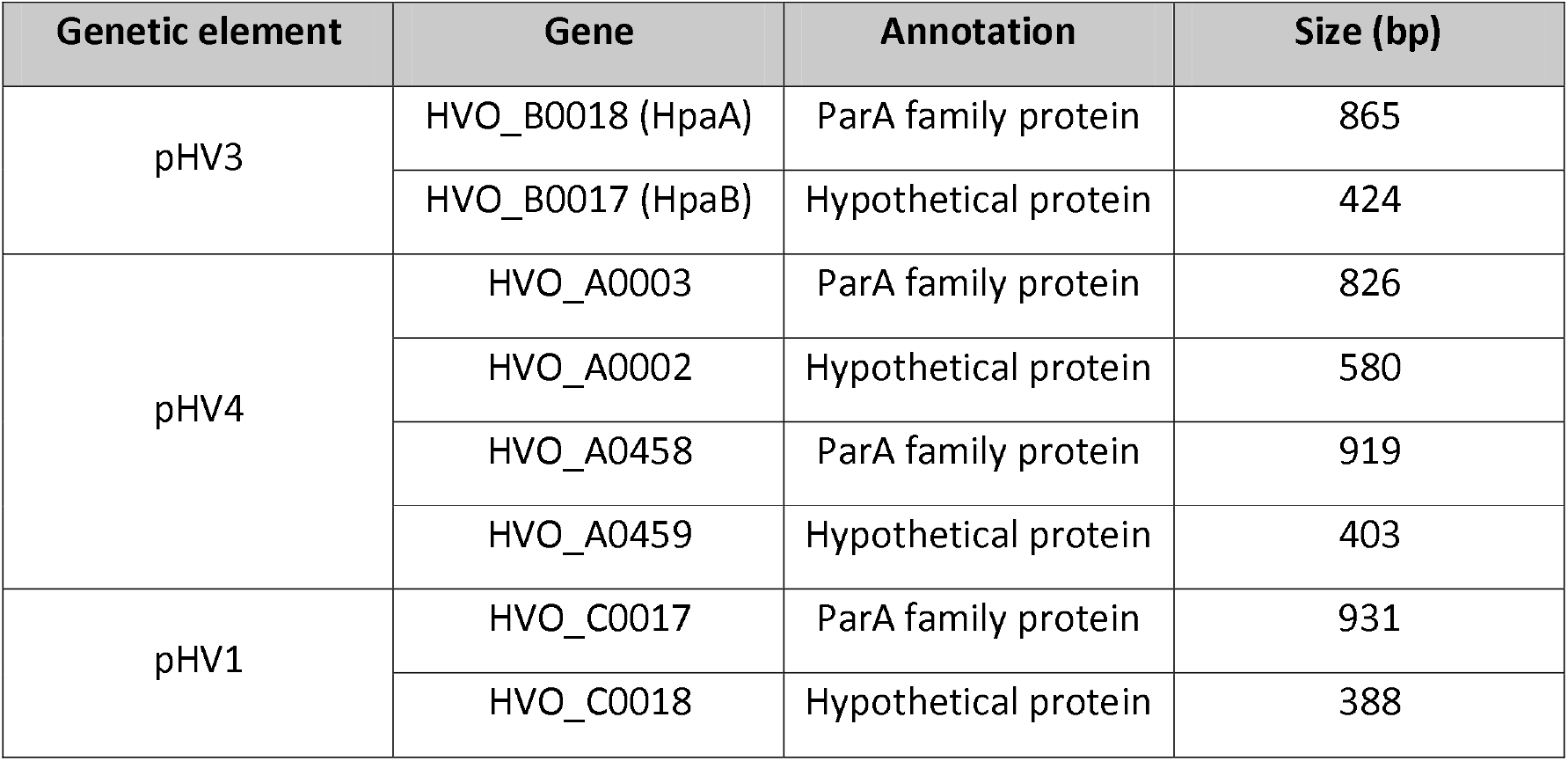
Four open reading frames on the *H. volcanii* genome with a ParA annotated protein.

| Genetic element | Gene | Annotation | Size (bp) |
| --- | --- | --- | --- |
| pHV3 | HVO_B0018 (HpaA) | ParA family protein | 865 |
|  | HVO_B0017 (HpaB) | Hypothetical protein | 424 |
| pHV4 | HVO_A0003 | ParA family protein | 826 |
|  | HVO_A0002 | Hypothetical protein | 580 |
|  | HVO_A0458 | ParA family protein | 919 |
|  | HVO_A0459 | Hypothetical protein | 403 |
| pHV1 | HVO_C0017 | ParA family protein | 931 |
|  | HVO_C0018 | Hypothetical protein | 388 |

In this project we focus on the secondary chromosome pHV3, with the operon containing the HVO_B0017 (*hpaA*) and the HVO_B0018 (*hpaB*) loci. The operon has an overlap of 4 nucleotides between the genes, and is therefore likely a single transcription unit.

The *hpaB* gene is located immediately downstream of *hpaA*, suggesting that the expressed protein might be a functional ParB homologue. However, upon searching the protein sequence of HpaB against the NCBI database using the Basic local alignment search tool (BLAST) no significant similar proteins were obtained. Further, HpaB showed no significant similarity to any annotated or characterized bacterial protein, including ParB. In contrast, the protein sequence of HpaA showed a high degree of similarity (70 % – 90 %) with other annotated ParA-like proteins in other Haloarchaea, and specifically *Haloferax sp*.

To assess the similarity between HpaA and canonical ParA proteins, a multiple sequence alignment was performed using the HpaA sequence and characterized ParA family proteins from *Escherichia coli, Bacillus subtilis, Corynebacterium glutamicum, Mycobacterium tuberculosis, and Clostridium Sp*. Additional comparisons included ParA-related proteins such as MinD, MipZ, and McdA from *Cyanothecae*, as well as SegA from the archaeon *Sulfolobus*. HpaA shows a high degree of similarity with the conserved ATPase Walker motifs of canonical ParA family proteins. This is corroborated by the presence of conserved key residues in the Walker A region, Walker A’ region, and Walker B region; especially the signature lysine, catalytic lysine, and invariant glycine in Walker A (Figure 1a).

A phylogenetic tree constructed using the neighbour joining method with the aforementioned protein sequences categorized the ParA family proteins into three distinct clusters (Figure 1b). The clusters were determined using phylogenetic distances dependent branch lengths. Clustering of similar proteins like MinDs, and canonical ParAs indicated the robustness of the tree. In the calculated tree, HpaA clustered with MipZ protein from *Caulobacter vibriodes*. The tree indicates a higher similarity of HpaA to MipZ and McdA due to branch closeness as compared to the similarity of HpaA with other ParA-like proteins like ParA, or MinD.

Comparison of the predicted structure of HpaA using Alphafold 3 with known structures of MipZ (pdb: 2XJ4) and ParA (pdb: 3EZ2) showed that HpaA shared a higher structural similarity with MipZ as compared to ParA (Figure 1c, 1d). The RSMD scores between the predicted HpaA structure and MipZ were lower at 9.931 Å unpruned and 0.93 pruned; while ParA from E. coli had higher RSMD scores at 10.416 Å unpruned and 0.99 Å pruned. However, in both cases, the conserved ATPase Walker motifs overlapped to a high degree. ParA-like proteins such as ParF, Soj, MipZ, and ParA form dimer sandwiches that trap nucleotides for hydrolysis, with the signature lysine of each monomer inserted into the active site of the other ^28,80-82^. McdA forms a similar dimer sandwich without the signature lysine and rather relies on an McdA-specific conserved lysine ^83^. On the other hand, SegA forms a “forward-backward” dimer rather than adopting an ATP sandwich ^84^. Structure prediction of dimeric HpaA in presence of ATP via Alphafold 3 shows ATP sandwiched within the dimer interface with the signature lysine (K11) predicted to interact with the ATP of the other subunit, similar to canonical ParAs described above (Supplementary figure 1a). Moreover, the surface of the predicted HpaA dimer is largely negative owing to the high salt concentration of *H. volcanii* cytoplasm. However, it contains a positively charged cleft (Supplementary figure 1b, c) which is predicted to be the DNA binding region.

### Dynamic localisation of HpaA depends on its putative conserved ATPase cycle residues

As fundamental insights toward the understanding of canonical ParABS systems can be collected via the study of their subcellular localization, we proceeded to tag HpaA with msfGFP followed by in trans expression in the H26 wild type, and in *H. volcanii* H26 *ΔhpaA*. HpaA was observed to form patchy foci in a compacted area towards the centre of the cells (Figure 2a arrows), and also form a cloudy dispersed population throughout the cells. In some cases, where foci in WT cells were absent, HpaA showed oscillatory behaviour (Figure 2b, Supplementary video 1). This was predominantly observed in rod shaped cells. Wild type HpaA oscillated in strains deleted for the native hpaA allele as well as in strains also expressing native *HpaA*, moving from one pole to the other in a short amount of time (∼5 minutes), and persisted longer at the poles.

**Figure 2.**
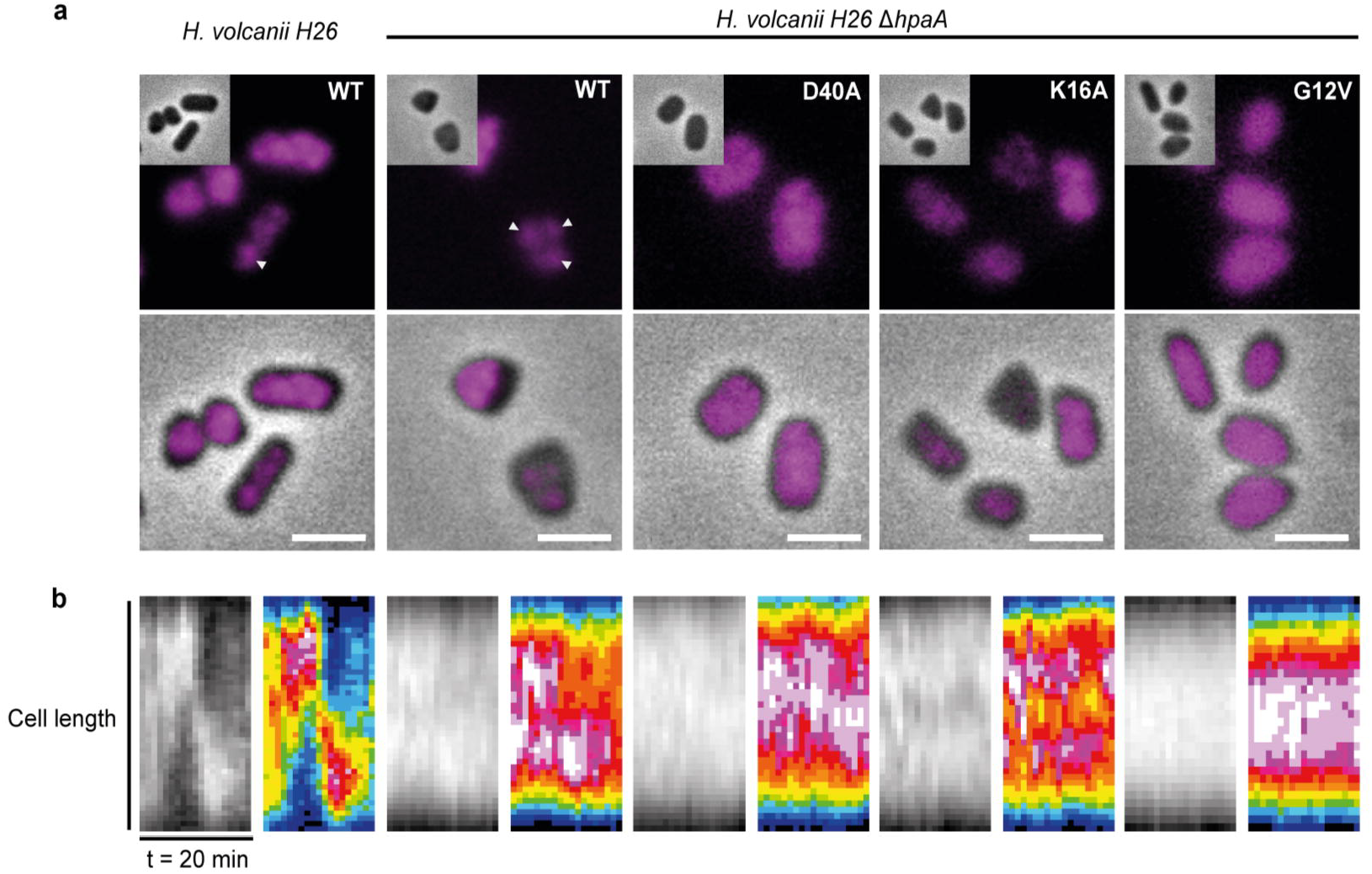
*In vivo* localization and dynamics of HpaA depends on key residues in the walker motif. (A) Fluorescence microscopy showing different localizations of variants of HpaA tagged with msfGFP (magenta) in wild type and *ΔhpaA H. volcanii* H26. Scale bar = 2 µm, arrows indicate foci. (B) Kymographs of representative cells which showed oscillation with wild type HpaA in both strains, and the lack thereof in mutants. Supplementary videos 1 and 2 show oscillation of HpaA-msfGFP in wild type cells, and lack thereof in cells lacking HpaB.

Subcellular dynamics of ParA-like proteins depend on their ATPase cycles. The Walker sites harbour highly conserved amino acid residues, which are found in both bacterial ParA proteins and HpaA (Figure 1a). These residues are essential for the ATP-dependent functions of dimer-sandwich forming ParA-like proteins ^81,85,86^. Any variations in these amino acids disrupt ATP binding, hydrolysis, or dimer formation, ultimately impairing the protein’s localization and function in vivo. Strategic substitutions were introduced in the conserved key residues of HpaA, that would lead to HpaA being trapped in different stages of its mechanistic cycle. The invariant glycine was replaced with a valine at position 12 (G12V), the catalytic lysine with alanine at position 16 (K16A) and the aspartate at position 40 with alanine (D40A). These mutations have been previously described for a variety of ParA-like proteins ^29,40,81,82^.

All HpaA mutants were expressed in a strain background lacking wild type HpaA to prevent heterodimer formation. None of the mutants showed any oscillatory behaviour. The kymographs instead indicate a more stable localization of the mutants (Figure 2b). The only mutant to show patchy localization *in vivo* was the putative ATP-binding mutant HpaA K16A (Figure 2a). On the other hand, HpaA-msfGFP D40A had a more diffuse signal with a higher concentration of the protein towards the center of the cell-consistent with a nucleoid bound protein, while HpaA-msfGFP G12V showed a near homogenous distribution of fluorescence. The difference in the localization of the different variants of HpaA indicate the importance of the conserved residues in its walker motifs, providing solid cell biological evidence that HpaA functions as ATPase just as bacterial ParA-like proteins.

### HpaA localisation and dynamics are dependent on HpaB

It is known that ParA-like protein dynamics, and in certain cases intracellular localization is dependent on the activity of ParB or non-homologous cargo adapters ^85,87-89^. We aimed to determine whether HpaA localization and dynamics also depend on presence of HpaB. To this end, *hpaA-msfGFP* and *hpaA-Halo* were ectopically expressed in *H. volcanii* H26 strains wild type and *H. volcanii* H26 *ΔhpaB*. HpaA-msfGFP was used to qualitatively describe the dynamics of HpaA, while the Halo tagged protein was constructed to study the effects of HpaB on HpaA on a single molecule level using organic dyes. HpaA and HpaB were also tagged with mTurqoise2 and mScarletI ^90^, respectively for dual fluorescent expression and co-localization studies.

Fluorescence images show that a majority of the wild type cells in logarithmic stage of growth display a heterogenous distribution of HpaA-msfGFP all over the cell. In most cases, HpaA additionally tends to form foci like structures (Figure 2; Figure 3a). On the other hand, in the absence of HpaA, the fluorescence of HpaA-msfGFP is homogenously distributed in the cells, and no discrete foci were observed (Figure 3a). Moreover, the oscillatory behaviour of HpaA seemed to be abolished in absence of HpaB (Supplementary Video 2). The plate-like cells showed a complete absence of oscillation of HpaA in both wild type and mutant strains. Upon expression of fluorescently tagged HpaA and HpaB, foci from both proteins were observed to often localize in close proximity (Figure 3b). The presence of foci with non-obvious colocalization can be attributed to the wild type background of the strain, expressing the native copies of both HpaA and HpaB. A representative line scan of foci from each channel (HpaA-mTq2 and HpaB-mSC-I) shows that there is enrichment of HpaB and HpaA around the other foci as well. Combined with the observation that HpaA foci formation requires the presence of HpaB, the colocalization between HpaA and HpaB supports a model in which HpaB causes a local enrichment of HpaA (Figure 3b).

**Figure 3.**
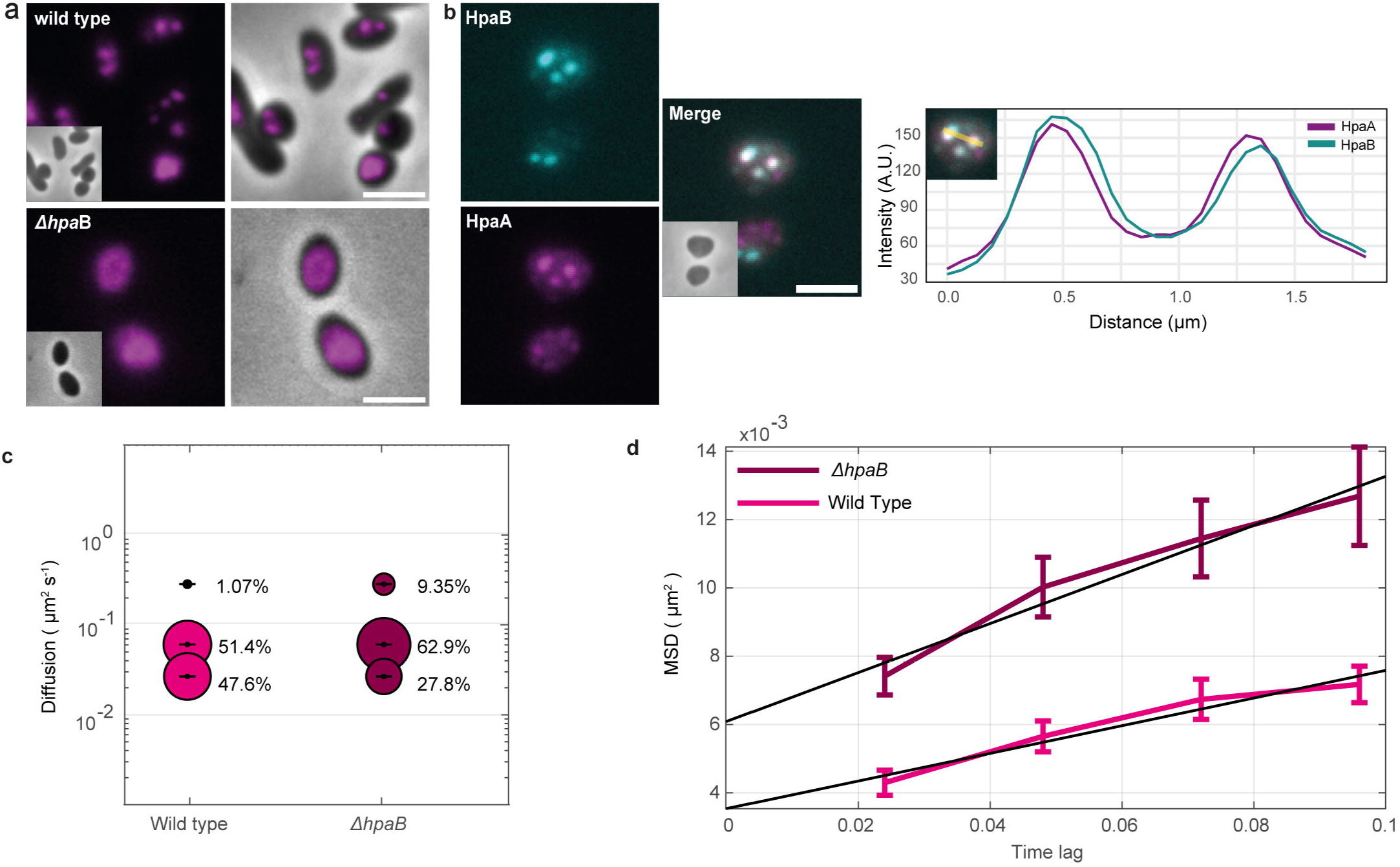
*In vivo* localization and dynamics of HpaA depend on HpaB. (A) Fluorescence microscopy showing representative cells with HpaA tagged with msfGFP (red hot intensity) in wild type cells and cells lacking *hpaB*. (B) Colocalization of HpaA-mTurquoise2 (magenta) and HpaB-mScarletI (Cyan) foci in representative cells along with a representative line scan showing the overlapping peaks of foci (yellow line in inset shows selected area of interest). Scale bar = 1 µm. (C) and (D) show dynamics of single particles of TMR stained HpaA-HaloTag in *H. volcanii* H26 *WT* and *ΔhpaB* using SMTracker (Rösch et al, 2018). (C) Bubble plot showing single-molecule diffusion rates of HpaA-HaloTag in wild type and mutant cells. Populations were determined by fitting the cumulative distribution function (CDF) of square displacements with a three-component model (‘fast’, ‘slow’, and ‘static ‘protein populations) (D) Plot of the mean-squared displacement of HpaA-Halo in wild type and mutant cells.

To further investigate the effect of HpaB on HpaA dynamics and localization, single-particle tracking (SPT) was performed using single-molecule localization microscopy (SMLM). Therefore, ectopically expressed HpaA-HaloTag was stained with the TMR fluorescent dye to enable precise single-molecule detection and tracking within live cells. SPT measurements used a 20 ⍰ ms exposure time.

Comparison of single molecule dynamics showed that HpaA was more mobile in absence of HpaB as compared to the wild type cells (Figure 3c, 3d). Protein dynamics are comprised of multiple separate subpopulations, each defined by its own parameters. To account for this behaviour of dynamic proteins, jump distance (JD) analysis was performed. The data could be interpreted best as three distinct populations which were identified and are represented as bubble plots (‘fast’, ‘slow ‘and ‘static’ from top to bottom) in both strains (Figure 3c). The fast-mobile proportion of the protein was small to begin with, at 1.07 %. However, the proportion of fast mobile protein in the *ΔhpaB* strain shows a huge increase to 9.35 %. Moreover, an increase in the slow mobile fraction of the protein can also be observed from 51.4 % in the wild type to 62.9 % in the mutant. The increase in both fast and slow mobile populations is mirrored by corresponding decrease in the static fraction from 47.6 % in wild type to 27.8 % in *ΔhpaB* cells (Figure 3c). In light of these results, we hypothesize the three fitted populations of the proteins to be enriched respectively in diffusive monomers (fast), DNA interacting dimers (slow diffusive), and the segregation complex associated dimers (static). The distribution of the populations based on their dynamics showed that the absence of HpaB led to an increased mobility of HpaA-HaloTag, indicating that HpaB might have a role in at least transiently stabilizing the HpaA interaction to the segregation complex. In the absence of HpaB, decreased recruitment of HpaA onto the segregation complex would lead to an accumulation of dimeric HpaA interacting non-specifically with DNA hence increasing the slow diffusive population and decreasing the static population. Subsequently, saturation of DNA with HpaA would lead to freely diffusive population of HpaA increasing, which can be seen as the highest increase in fast diffusive population. Mean-squared displacement (MSD) was used as a quantitative indicator of the averaged diffusion speed. It showed that the diffusion coefficient of HpaA-HaloTag was larger in absence of HpaB (Figure 3d). The results are intriguing because HpaA loses its oscillatory behaviour, while its dynamicity increases in absence of HpaB. This is probably because of an increase in free cytoplasmic protein, and by extension an increase in random protein diffusion.

### HpaB is essential to maintain the secondary chromosome pHV3

Chromosomes and extra-chromosomal elements like plasmids are self-replicating entities allowing their vertical inheritance across generations. Inheritance of genomes via ParAB like systems involves and requires specificity, and would need to code for the segregation complex. Organisms with multiple chromosomes such as *Vibrio spp*, and *Deinococcus radiodurans* have *parAB* operons and distinct *parS* sites on their different chromosomes ^91,92^. Similarly, low copy number plasmids in bacteria, like the F plasmid or the P1 plasmid in E. coli can be stably maintained with the help of partition systems that are responsible for positioning them inside the cell ^93^. In contrast, high copy number plasmids usually do not need an active segregation system because random segregation is sufficient.

To test whether maintenance of pHV3 is affected upon deletion of *hpaA* or *hpaB*, we performed a qPCR to obtain a ratio of the secondary chromosome to the main chromosome in a given population. As under physiological conditions, the pHV3 to chromosome ratio ranges between 1.0 - 1.2, a significant decrease in this ratio upon deletion of the functional HpaAB system would indicate an overall decrease of pHV3 numbers in the population due to a loss of maintenance or stable inheritance.

Multiple biological replicates were performed with cultures of the *H. volcanii* wild type, *ΔhpaB*, and *ΔhpaA* strains to asses pHV3 to chromosome ratios. The deletion of hpaB resulted in the most significant decrease in the pHV3 to chromosome ratio (Figure 4a), from a mean ratio of ∼ 1 in the wild type to ∼0.35 in the ΔhpaB mutant. In contrast, the deletion of hpaA showed no significant change with the mean ratio staying at ∼0.9, indicating that HpaB is essential in maintaining the secondary chromosome to main chromosome ratio. Furthermore, we constructed complementation strains in which *hpaB* and *hpaA* were expressed from a plasmid under control of a tryptophan inducible promoter respectively. Synthesis of HpaA or HpaB was induced with 0.1 mM tryptophan. While over-expression of HpaA also shows a mild increase in the pHV3 ratio; ∼1.15, complementing HpaB significantly rescued the pHV3 ratio by a factor of ∼1.8, as the mean secondary chromosome to main chromosome ratio changes from ∼0.35 to ∼0.65.

**Figure 4.**
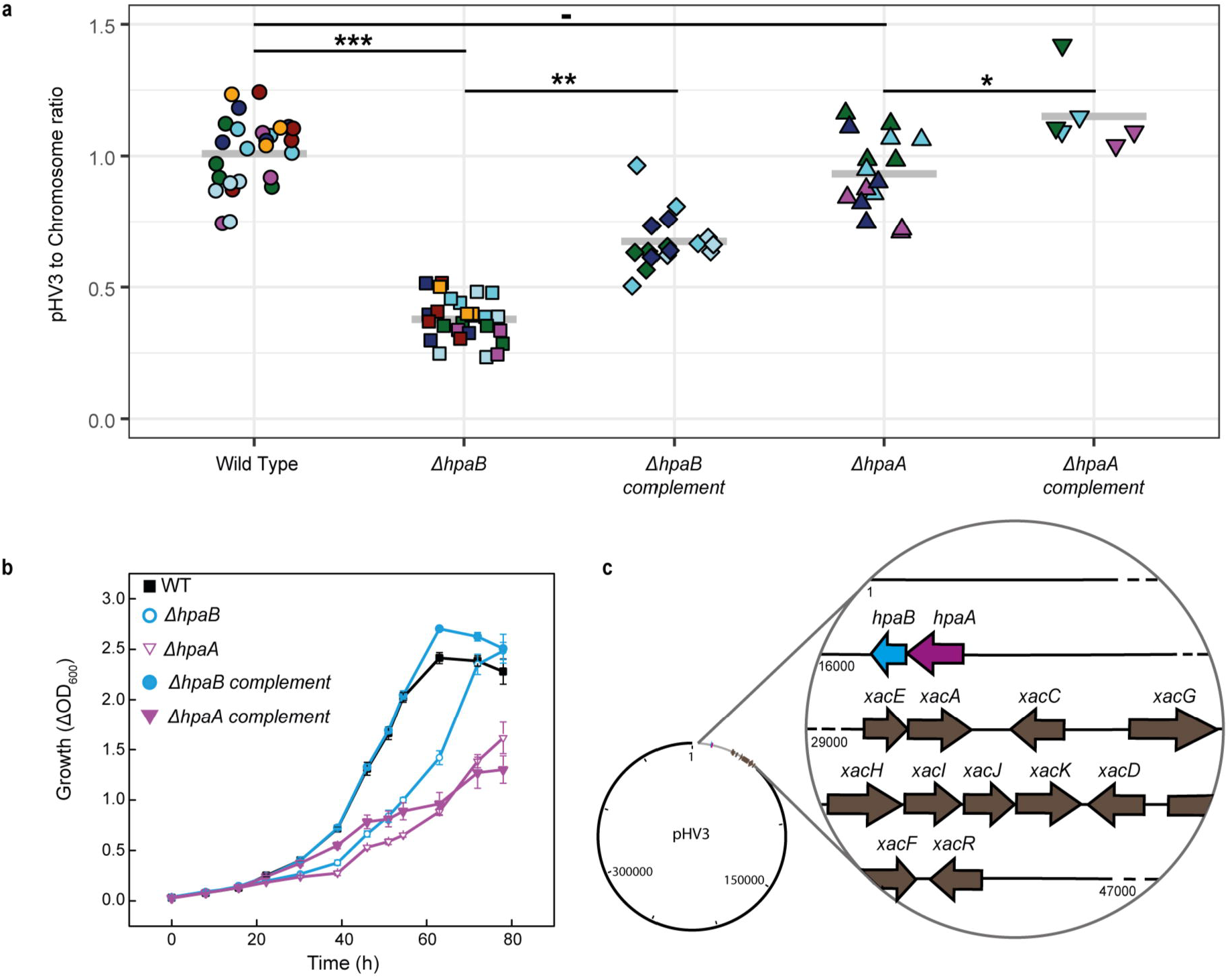
HpaB is essential to maintain pHV3 to chromosome ratio, and for growth on xylose. (A) qPCR assays performed on genomic DNA of *Haloferax volcanni* with primers for the main chromosome and pHV3 show a decrease in pHV3 to chromosome ratio (y-axis) for the mutants (x-axis) without HpaB. This decrease can be partially complemented by over-expression of the protein. Strains used: H26 *ΔhpaB, ΔhpaA*, wild type, and compliment strains induced with 0.1 mM tryptophan. Colours indicate biological replicates - in sequence: dark green, cyan, magenta, dark blue, dark red, light blue, and orange each with 2-4 technical replicates. Stars indicate significance: *** (<0.005), *(<0.5). (B) Growth curves of *H. volcanii* H26 wild type, *ΔhpaB, ΔhpaA*, and complement strains on synthetic media containing 20 mM xylose showing a significant defect of growth in the mutants which is rescued by complementation. All curves were performed in four replicates, error bars depict standard deviation. (C) Cartoon showing the position of the hpaAB operon and the xylose degradation pathway genes on pHV3.

### HpaA and HpaB are essential for growth on xylose

A fundamental aspect of chromosomes is that they carry housekeeping and core genes that are usually indispensable for viability and cell fitness. This includes genes involved in uptake and metabolism of carbon sources. The secondary chromosome pHV3 contains all genes that encode proteins involved in xylose uptake and degradation (Fig. 4c). In *H. volcanii*, D-xylose is metabolized via an oxidative degradation pathway to α-ketoglutarate ^94,95^. The corresponding enzymes are encoded by the xac gene cluster, encoding xylose dehydrogenase (XacA, HVO_B0028), pentonolactonase (XacC, HVO_B0030), D-xylonate dehydratase (XacD, HVO_B0038A), 2-keto-3-deoxy-xylonate dehydratase (XacE, HVO_B0027), and α-ketoglutarate semialdehyde dehydrogenase (XacF, HVO_B0039). Within the same cluster, the genes *xacGHIJK* (HVO_B0034–HVO_B0038) encode an ABC transporter for the uptake of xylose, consisting of a substrate-binding protein (XacG), two transmembrane domains (XacH and XacI), and two nucleotide-binding domains (XacJ and XacK) ^96^. The expression of both the catabolic and transporter genes is controlled by the transcriptional activator XacR (HVO_B0040), which induces their transcription in the presence of xylose and arabinose ^97^.

We reasoned that the secondary chromosome pHV3 may become in particular important during growth on xylose as a sole carbon source. Consequently, a decrease in pHV3 to chromosome ratio due to loss of inheritance via mis-segregation would lead to a defect in growth on xylose. In line with our hypothesis, deletion of *hpaB* resulted in severe growth defects. Moreover, despite not affecting the pHV3 to chromosome ratio, the deletion of *hpaA* also resulted in growth defects, indicating that both genes are essential for robust growth under these conditions (Figure 4b). Moreover, these defects could be rescued upon tryptophan-controlled expression of HpaA and HpaB via ectopic complementation constructs. Complementation of *hpaB* fully restored growth to wild-type levels. In contrast, complementation of *hpaA* initially improved growth relative to the deletion strain, but growth plateaued over time, and prolonged cultivation failed to achieve full recovery. The transient rescue observed upon ectopic expression likely reflects copy-number sensitivity of the HpaAB system, in which sustained complementation is not achieved because proper function requires balanced stoichiometry with interacting partner proteins. This is generally difficult to achieve with plasmid-born inducible expression systems.

These results indicate that the concentration of HpaA in the cell might be important for full functionality and *hpaA* expression likely requires precise temporal or spatial regulation, for full function during xylose-dependent growth. In contrast to growth on xylose, HpaA or HpaB are not essential for viability of *H. volcanii* H26 when grown on glucose as a carbon source (Supplementary Figure 2).

### HpaB is a putative DNA-binding protein localizing in foci

Most ParAB-like systems involved in cargo positioning exist in operons containing an adapter protein ^98,99^. This adapter binds to the cargo; for example, centromere binding proteins (CBPs) like ParB binds to the *parS/parC* sites on chromosomes or plasmids ^100,101^, and McdB binds to carboxysomes ^38,99,102^. CBPs are known to form condensed structures to allow high concentration of the protein around their cargo sites ^30,102-104^.

We hypothesize HpaB to be the putative DNA binding adapter for the HpaAB system, since it is encoded immediately downstream of HpaA, is a relatively small protein, and the open reading frame contains no other protein with a DNA-binding helix-turn-helix (HTH) or ribbon-helix-helix (RHH) motif. Though HpaB showed no sequence homology or significant similarity with any known CBP or analogous proteins, structure alignment of predicted structures showed the presence of similar domains with other CBPs like ParG^105^, and SegB^84^, particularly in its RHH motif. ParG (PDB: 1P94)^105^ in particular showed a similar RHH fold with an overall predicted RSMD of 1.0 Å (Supplementary figure 3a). On the other hand, when the predicted structure of HpaB was compared with the known structure of SegB (PDB: 7DUV)^84^, an RSMD score of 0.525 Å was predicted with higher similarity in the monomers rather than the dimer (Supplementary figure 3b). Furthermore, structure prediction via Alphafold 3 indicated a presence of the DNA binding region within its RHH motif (Figure 5a). Two residues, histidine at position 70 and tyrosine at position 74, were predicted to interact with DNA. To test whether these residues had any influence on the in vivo localization of HpaB, variants were generated with both residues mutated to alanine to cause a loss of function. In vivo localization was studied by expressing fluorescent tagged HpaB. Wild type HpaB-msfGFP showed presence of clear foci in the cells (Figure 5b). While HpaB H70A was able to form foci, HpaB Y74A and a double mutation lacked any discernible foci. The results indicate that the foci formation of HpaB depends on DNA binding via the tyrosine residue at position 74 in the RHH motif.

**Figure 5.**
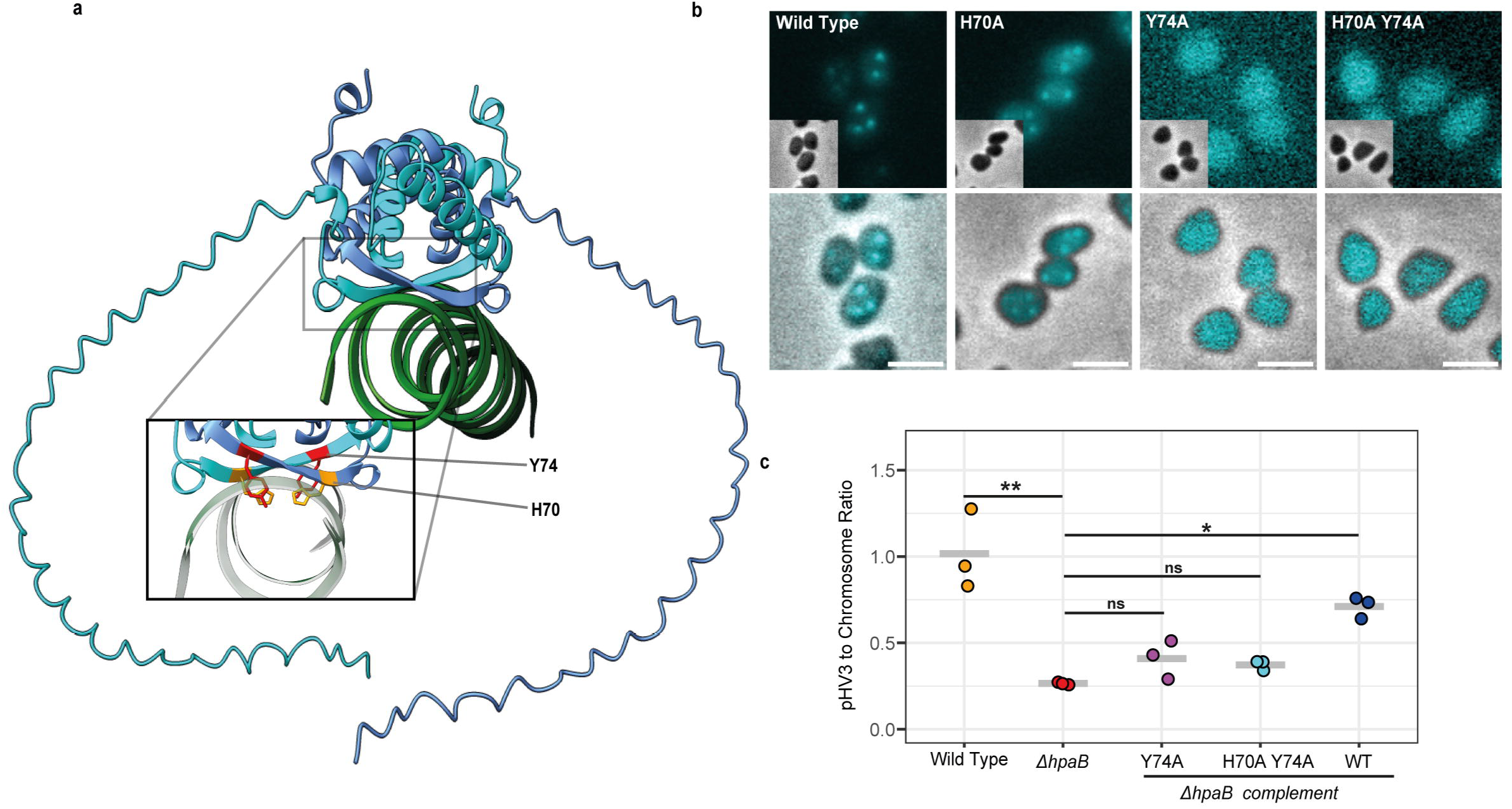
HpaB requires tyrosine at position 74 to form foci. (A) Predicted binding of HpaB dimer with DNA, highlighting important residues in the ribbon-helix-helix motif that interact with the DNA strand, tyrosine (Y) in red and histidine (H) in orange. (B) Widefield fluorescence microscopy showing representative cells with fluorescence of HpaB variants tagged with msfGFP in wild type cells showing foci and point mutation variants lacking foci. Scale bar = 1 µm. (C) qPCR assays performed on genomic DNA of *H. volcanii* showing the partial complementation of pHV3 to chromosome ratio by over-expression of WT HpaB, and no complementation by HpaB variants. Stars indicate significance: *** (<0.005), *(<0.5).

We further tested via qPCR whether the foci formation phenotypes resulted by the mutation of these residues is associated with defective pHV3 maintenance. The strain lacking HpaB was complemented with ectopic expression of wild type HpaB, HpaB Y74A, and HpaB H70A Y74A. While the wild type HpaB is able to partially complement the pHV3 to chromosome ratio from ∼0.35 to ∼0.65, the other two variants show no significant complementation with the pHV3 to main chromosome ratio in both HpaB Y74A, and HpaB H70A Y74A being around 0.4. These results indicate a high level of correlation between the role of HpaB in maintaining the secondary chromosome in the population, and its ability to form foci; both of which rely on the tyrosine at position 74.

### HpaB foci distribute homogeneously in an HpaA dependent manner

Since HpaB is an uncharacterized hypothetical protein with structural homology to proteins like ParG and SegB, it was important to determine the behaviour of the protein *in vivo*. The distribution of HpaB in presence and absence of HpaA would give insights on the role of HpaB. Thus, HpaB-msfGFP was ectopically expressed in *H. volcanii* H26 wild type and *ΔhpaA* cells. Microscopic examination of HpaB-msfGFP showed the presence of HpaB condensed in foci (Figure 6a). *H. volcanii* cells contained 2-3 foci per cell in presence of natively expressed HpaA. Deletion of *hpaA* led to a significantly increased average number of foci ∼ 8 foci per cell (Figure 6b). Moreover, the HpaB foci seemed to be closer together in absence of HpaA. The high density of the foci in the *ΔhpaA* mutant can be attributed to the fact that the foci number shows a 2-3-fold increase, and are likely not able to fuse without HpaA.

**Figure 6.**
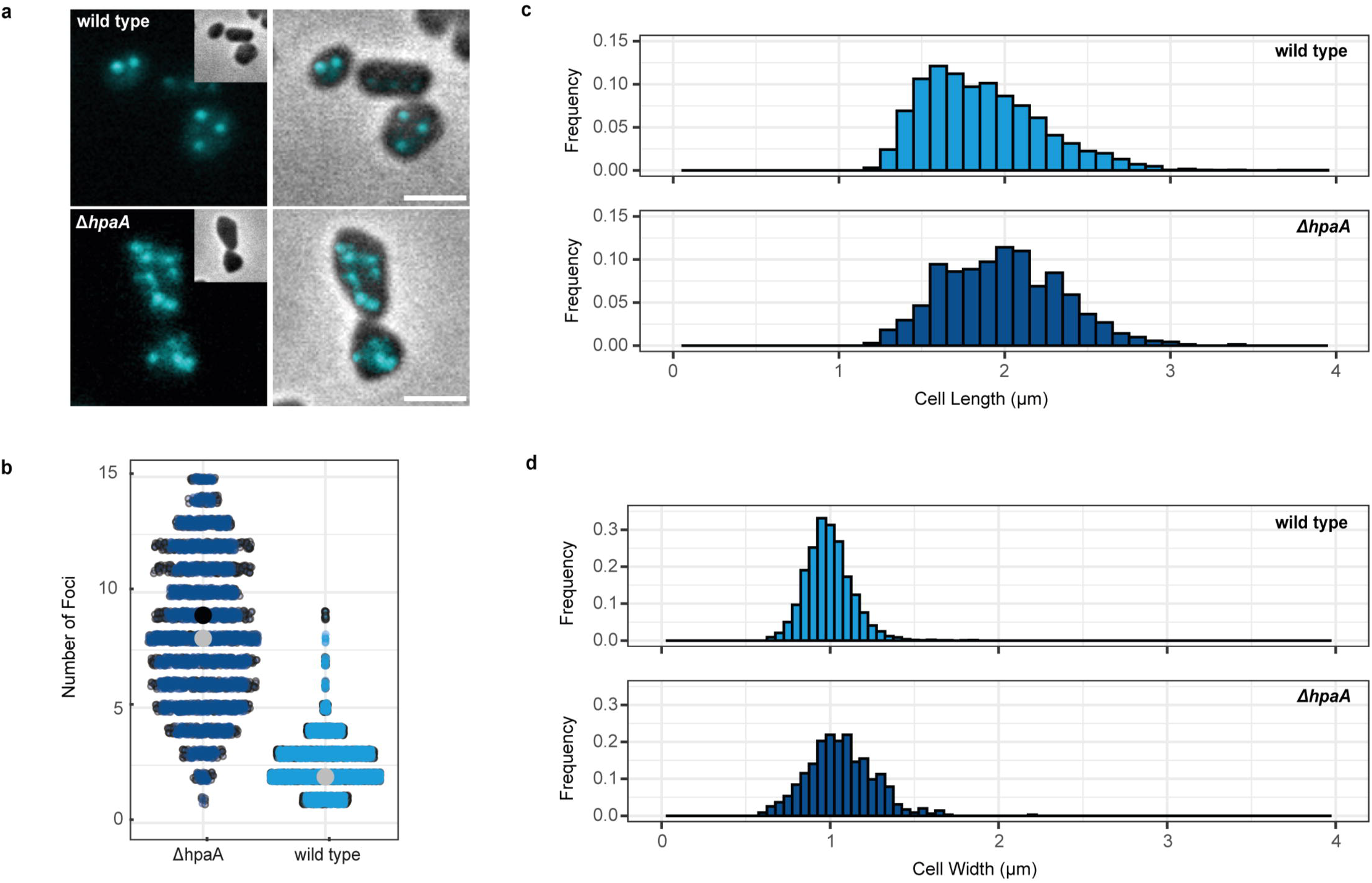
HpaB foci numbers are dependent on HpaA. (A) Widefield fluorescence microscopy showing representative cells with fluorescent foci of HpaB tagged with msfGFP (cyan) in wild type cells and cells lacking *hpaA*. Scale bar = 1 µm. (B) Comparison of number of foci per cell (y-axis) in wild type and mutant (x-axis); p value < 1e-16, n > 7000 cells (>3500 cells per replicate). Colours for each strain indicate two biological replicates. (C) and (D) show how the deletion of hpaA affect the dimensions of the cells, with the mutant cells showing a non-significant increase in length and width.

To compare the distribution of HpaB foci quantitatively, the microscopic data of wild type and *ΔhpaA* cells, along with the distribution of fluorescence was extracted as X and Y data points. These were plotted together to obtain the distribution of foci in an average cell. The nearest neighbour distribution for each focus within a cell shows that HpaB foci are significantly closer to each other in absence of HpaA (Figure 7a), as indicated by left shift on the cumulative nearest distance (Figure 7b). Moreover, HpaB foci are differently distributed in the presence and absence of HpaA, which a lack of HpaA causing a shift from ordered to a random distribution. The heatmap (Figure 7c) shows visually where HpaB foci are located, and it is clear that HpaA is required for an ordered distribution of the foci such that they neither cluster in the centre of the cell, nor near the membrane. These data clearly support a model of an active distribution of HpaB foci by the HpaA ATPase via corralling a few HpaB-DNA complexes together. The number of foci in the absence of HpaA (averaging at around 8) tends to be closer to the number of replicons, due to decreased corralling. This indicates that pHV3 copies possibly need corralling via HpaA to be bundled together into HpaB foci.

**Figure 7.**
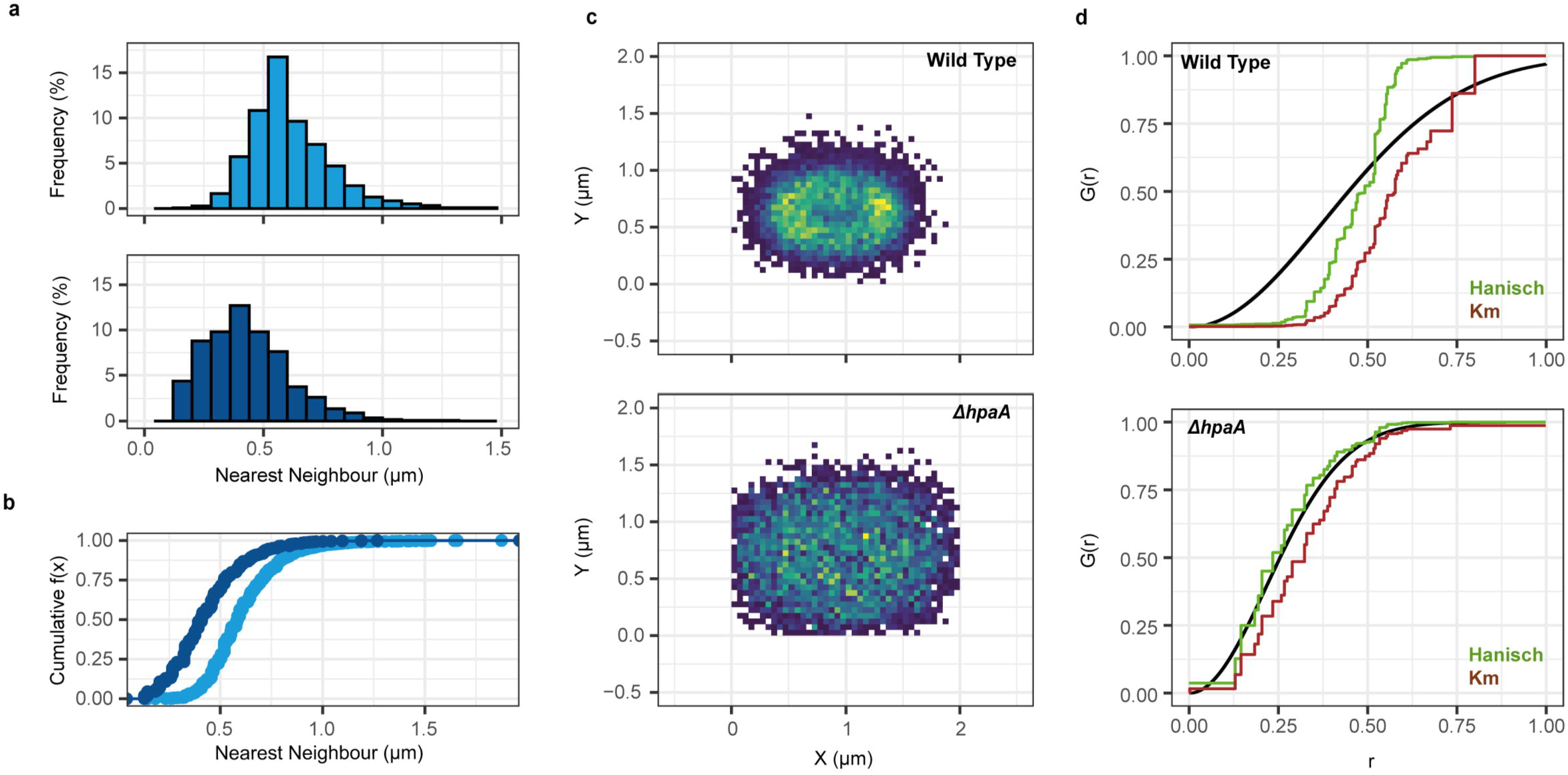
Distribution of HpaB foci is HpaA dependent. (A) The histogram distribution of the distance between closest foci within each cell (x-axis) and the frequency in which they are observed (y-axis). Cyan is wild type, dark cyan is mutant (B) Cumulative nearest neighbour distances showing a left shift in the mutant strain. (C) The distribution heatmap of foci within all the analysed cells centred using X/Y coordinates showing an average likelihood of foci distribution in cells. (D) G-function curve with border corrected nearest neighbour cumulative curves using Hanisch (green) and km estimates (brown). The black curve indicates theoretical randomness on the given set of data; the corrected curves show that the HpaB foci are non-randomly distributed in an HpaA dependent manner. p value < 1e-16, n > 7000 cells.

To quantify the randomness, a G function curve was plotted for both the strains using the cumulative nearest neighbour distance between the HpaB foci within each strain. The G function G(r) measures the distribution of nearest distances of points (foci in our case). Given any set of points on a plane, random distribution of the points would lead to clustering and repelling behaviour within the same plane, while ordered distribution would tend closer to equipositioning of the points on the plane (Supplementary Figure 4). This distribution is then plotted as a G function curve with the cumulative nearest neighbour distances (ecdf) for each point, and corrected for border effects due to a limited plane size using the Hanisch and Kaplan-Meier-estimates ^106,107^ (Supplementary Figure 4). In case of random distribution, the corrected curves tend closer to complete spatial randomness (CSR), while an ordered distribution would tend towards compression of the distance on the x-axis which can be attributed to equal distances between each point (Supplementary Figure 4).

In our case the ecdf curves were plotted against a theoretical random distribution according to the provided dataset, and are then corrected via Hanisch and Km estimates^106,107^. A random distribution of the foci can be observed in the strain with *hpaA* deletion (Figure 7D), with the Hanisch and Km estimates tending towards the CSR. On the other hand, in the wild type cells, it can be seen that the foci start repelling each other while interestingly forming clusters at higher distances and G(r) values (Figure 7D). The G function curve tends towards a more equipositioning via ordered distribution of the foci with a preference of distance between foci being around 0.4 to 0.7 µm. These results allow us to conclude that HpaA is required for ordered distribution of the HpaB foci via equipositioning them in two dimensions, and its lack thereof leads to a random distribution of the HpaB foci.

### HpaB interacts with HpaA with an N-terminal domain

The disordered distribution of HpaB foci in the absence of HpaA suggests that the interaction between HpaB and HpaA is essential for *in vivo* localization of HpaB. Despite the diversity of cargo adapter protein in ParA-like operons, it has been shown that they often interact with their ATPase partner via the N-terminus ^89,108-110^.

To determine if HpaB might also have an N-terminus dependent interaction with HpaA; we modelled the predicted interaction between the two proteins via Alphafold 3 ^77^. The prediction included an ATP-dependent HpaA sandwich dimer, DNA, and two variants of HpaA: Wild type, and a truncated HpaB lacking residues 2-12 (HpaB_ΔN_) (Figure 8a). Both variants of HpaB were predicted to dimerize and showed DNA interaction via the RHH motif. The N-terminus of each monomer of HpaB formed a small α-helix comprising of 12 residues (Figure 8a inset), which was predicted to interact with a monomer of HpaA. Upon repeating this analysis without the 11 residues in the N-terminus of HpaB the α-helix did not form, and the proteins lost their interaction interface (Figure 8a).

**Figure 8.**
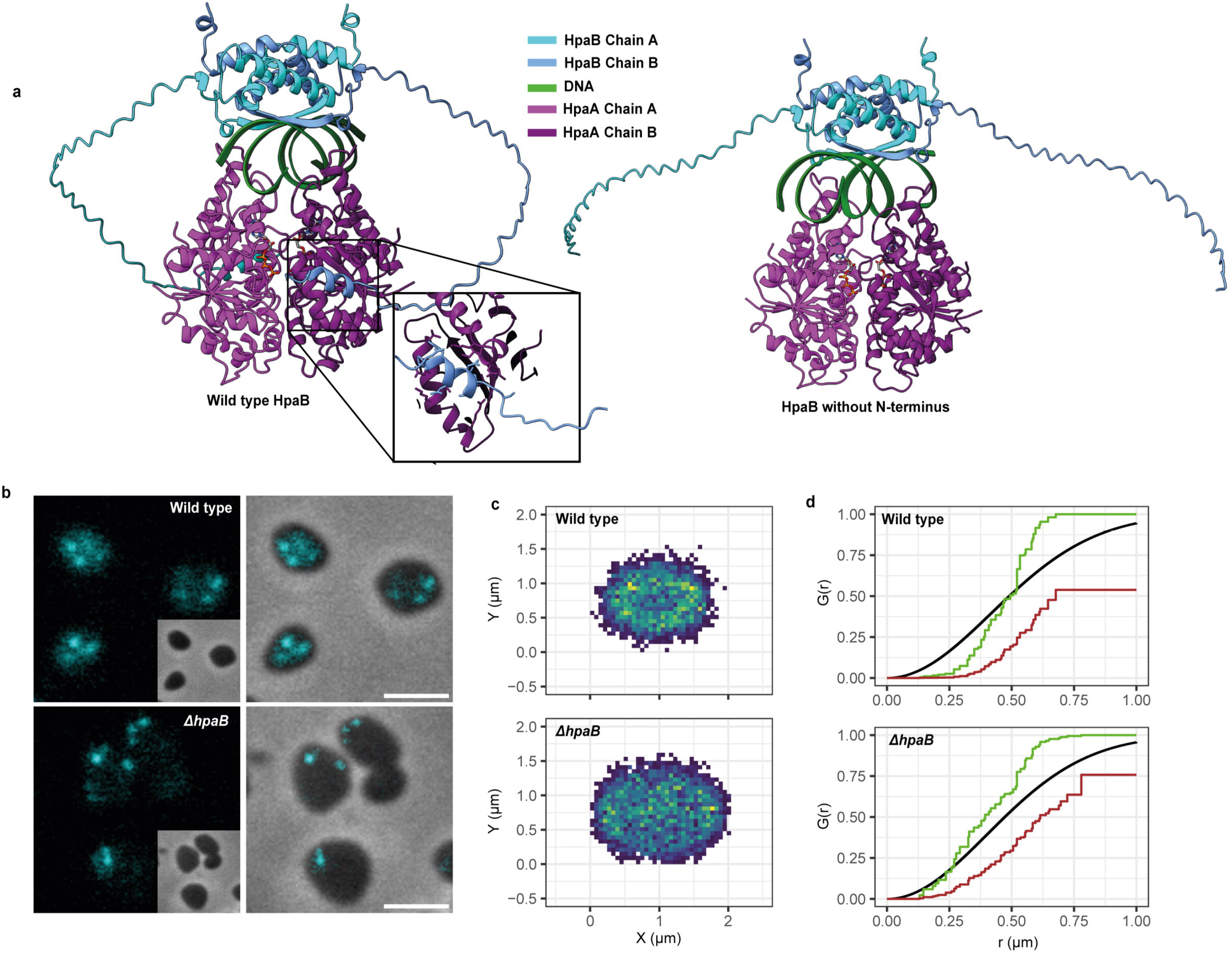
HpaB distribution depends on the presence of its N-terminus. (A) The predicted binding of HpaB_WT_ and HpaB_ΔN_ with HpaA dimers via the N-terminus of HpaB in focus (B) Widefield fluorescence microscopy showing representative cells with fluorescent foci of HpaB_ΔN_ tagged with msfGFP (cyan) in wild type cells and cells lacking *hpaB*. Scale bar = 1 µm. (C) The distribution heatmap of foci within all the analysed cells centred using X/Y coordinates showing an average likelihood of foci distribution in cells. (D) G-function curve with border corrected nearest neighbour cumulative curves using Hanisch (green) and km estimates (brown). The black curve indicates theoretical randomness on the given set of data; the corrected curves show that the HpaB foci are non-randomly distributed in an N-terminus dependent manner. p value < 1e-16, n > 3000 cells.

In order to corroborate these models, the *in vivo* role of the N-terminus was determined by expression of HpaB_ΔN_-msfGFP in wild type and *ΔhpaB H. volcanii*. The wild type cells contained both the native copy of HpaB and the HpaB_ΔN_-msfGFP, while the *ΔhpaB* strain expressed only the truncated HpaB. The N-terminus was found to not be essential for the foci formation behaviour of HpaB, as HpaB_ΔN_-msfGFP formed foci in both strains (Figure 8d). However, the localization of HpaB_ΔN_ revealed severe differences in both cases. Therefore, the localization and distribution were quantified in both strains as above. In the strain containing both native wild type HpaB and ectopically expressed HpaB_ΔN_, where heterodimers of HpaB and HpaB_ΔN_ might form, the foci showed a similar ordered distribution as wild type HpaB around the centre of the cell (Figure 8c), indicative of correct interaction with HpaA, which was shown before to be essential for correct HpaB positioning (Figures 6-7). On the other hand, the foci formed by homodimers of HpaB_ΔN_ lost this ordered distribution (Figure 8c) and instead had a random distribution as demonstrated by the ecdf curves (Figure 8d green). The random distribution of HpaB_ΔN_ is also apparent from the heatmap analysis (Figure 8c), mimicking those distributions of HpaB in the absence of HpaA (Figure 7). Therefore, we suggest from these results that the N-terminus of HpaB is highly likely the interaction domain with HpaA and therefore essential for the ordered distribution of HpaB foci.

## Discussion

### HpaA is a ParA orthologue

Bioinformatic analysis of HpaA shows a high degree of sequence similarity in the conserved ATP walker motifs with canonical bacterial ParA-ATPases, especially with respect to conserved catalytic and signature lysines, and the invariant glycine residues ^26,82^. In canonical ParA systems, dimerization and ATP binding are tightly linked to DNA interaction, enabling dynamic gradients for positioning cellular cargo such as plasmids ^80^. The ATP-binding region in the predicted structure of HpaA appears to be sandwiched between two monomers, similar to canonical ParA dimers, meaning that the ATP is stabilized by the signature lysine of the respective other monomer. This brings the P-loops and switch II motifs of both monomers in close proximity once the dimer is formed ^82^.

Alphafold3 predictions also suggest that HpaA forms an ATP sandwiched dimer as seen in Soj and MinD from *Bacillus subtilis* ^82,111^ for example. Interestingly, the phylogenetic analysis of the full length HpaA sequence places it closer to ParA-like proteins such as MipZ and McdA, rather than Soj or plasmid born ParA’s (Figure 1). The structure of the carboxysome positioning ATPase McdA contains an unusual dimer formation ^83^. In ATP bound McdA a lysine residue located far from the P loop facilitates ATP dependent formation of a nucleotide sandwich dimer that positions the two McdA monomers in opposite orientation. This non-canonical lysine is conserved among McdA homologs ^83,85^. The sequence of HpaA however, contains the canonical signature lysine in the walker motif (Fig. 1a), assuming that the dimerization is not McdA-like. On the other hand, the predicted binding site of HpaA with HpaB seems to be on each monomer rather than a cleft on the dimer interface (Figure 8a) arguing towards homology to MipZ for adaptor binding as MipZ also lacks the hydrophobic cleft seen in proteins like Soj ^81,82^.

The HpaA dimer forms a positively charged DNA-binding surface clustered with lysines and arginines that is embedded within an otherwise negatively charged protein surface which is likely an adaptation to the hypersaline cytosol of *H. volcanii*, which contains close to 2 M KCl (Supplementary Figure 1). Since HpaA lacks a known DNA-binding motif, the positively charged patch is consistent with non-specific DNA interaction which is a hallmark of ParA-like ATPases ^82^. *In vivo* data support that HpaA behaves as a bona fide ParA-like protein. Walker motif mutants, including the putative ATP-binding mutant (K16A), ATP-hydrolysis mutant (D40A), and the invariant glycine mutant (G12V) all show localization patterns consistent with loss of ATP binding or dimerization, leading to either static foci or cytoplasmic dispersion, thereby phenocopying the subcellular localization known ParA-like proteins such as MipZ and McdA ^40,81,85,112^.

Interestingly, the G12V variant of HpaA exhibits homogenous cytoplasmic distribution, recapitulating the dynamics of equivalent McdA mutants ^85^. Furthermore, oscillatory behaviour of wild-type HpaA was observed in a subset of rod-shaped cells, suggestive of gradient-based positioning mechanisms similar to McdA ^38^, MinD ^39^, and ParA ^113^. The oscillation of HpaA was absent in cells lacking HpaB, suggesting that HpaB modulates HpaA localization, just like the adapter protein is responsible for modulating dynamics of the ParA-like ATPase in other ParAB-like systems ^85,88^.

### HpaB is a focus-forming, novel archaeal protein

HpaB was previously annotated as a hypothetical protein due to lack of functional characterization, but our data points towards it being a novel DNA-binding protein involved in chromosome partitioning. Sequence analysis shows homology to uncharacterized archaeal proteins but no similarity to bacterial homologs. HpaB is small (∼150 amino acids), disordered, and predicted to contain a ribbon-helix-helix (RHH) motif that are commonly found in centromere-binding proteins (CBPs) such as ParG ^105^ and SegB ^42^. The strongest evidence for DNA binding by HpaB stems from the Y74A mutant: substitution of the predicted DNA-contacting tyrosine residue abolishes focus formation, suggesting that the wild-type protein forms foci through direct interaction with DNA. Moreover, the qPCR data shows that the Y74A mutant is also unable to complement via relieving the secondary chromosome pHV3 to chromosome ratio in cells lacking HpaB. We interpret the observed foci to represent pHV3 clusters with bound HpaB. Such clustering mirrors known behaviours of ParB-like proteins that undergo liquid-liquid phase separation (LLPS) ^24^. The high salt environment in *H. volcanii* may promote or regulate LLPS, enabling HpaB to form localized condensates similar to McdB ^114^ or ParB ^24^. Currently, direct evidence for HpaB-pHV3 binding, and the specific site it binds to is missing and requires further experimental proof. However, the data obtained with the putative DNA-binding mutant strongly suggest that HpaB is directly binding its DNA cargo. Thus, we assume that HpaB is functionally analogous to the CBPs such as ParG from bacteria ^105^, or SegB from *S. solfataricus* that also bind DNA via its RHH motif and not via an adapter protein as seen in other archaeal systems such as the *Sulfolobus* NOB8H2 AspA-ParB-ParA system ^43^.

HpaB foci exhibit ordered spacing within cells, a phenomenon that is lost in the absence of HpaA. In analogy to the McdAB system ^38^, HpaA may prevent aggregation of HpaB foci, ensuring their equipositioning. Deletion of *hpaA* results in random distribution of HpaB foci, while deletion of *hpaB* alters the dynamics of HpaA, consistent with mutual interdependence. Moreover, this ordered distribution of HpaB is dependent on the presence of its N-terminus, which is predicted to interact with HpaA. Truncation of the N-terminus leads to a disordered distribution of HpaB foci, quite similar to the loss of ordered distribution in absence of HpaA. While the randomness is not as extreme, our data suggests that HpaB depends on an α-helix at the N-terminus to interact with HpaA; quite similar to how the N-terminus in known cargo adaptor proteins binds to their ParA-like partners ^86,110^. The differences in the foci number and the degree of randomness between the *in vivo* behaviour of truncated HpaB in absence of wild type HpaB, and behaviour of HpaB in absence of HpaA indicates a role of the N-terminus in more than just HpaA interaction. We speculate that the N-terminus of HpaB may be important for a more regulated condensation of HpaB via interaction with HpaA. Truncation of the N-terminus therefore leads to a more disordered and aggregating behaviour with the aggregated being randomly distributed in the cell.

### HpaA and HpaB form a hybrid partitioning system in Haloferax volcanii

The secondary chromosome pHV3 encodes genes involved in xylose metabolism ^96,97^. We therefore hypothesized that defects in chromosome segregation and unequal inheritance would impair growth and fitness when cells are cultivated with xylose as the sole carbon source. Consistent with this assumption, a pronounced growth defect was observed in defined medium containing xylose. The growth deficiency in the *ΔhpaB* strain was fully complemented by expression of *hpaB* in trans (Figure 4b). These findings support the conclusion that the absence of HpaB causes a segregation defect of chromosome pHV3. Inefficient segregation results in daughter cells that fail to inherit the secondary chromosome, leading to reduced overall population growth despite the survival of some cells retaining pHV3. Similar results were observed for the *ΔhpaA* strain; however, complementation in trans only partially restored growth (Figure 4b). Although initial growth was improved, the *hpaA* complementation strains eventually stalled. This incomplete rescue may reflect partial functionality of HpaA when provided in trans, suboptimal expression conditions, or the dynamic, concentration-dependent behaviour of HpaA, as reported for other pattern-forming or concentration-dependent proteins^115^. Further evidence for active partitioning of the secondary chromosome pHV3 comes from qPCR analysis, which revealed a significant reduction in the pHV3-to-chromosome ratio in *ΔhpaB* cells (Figure 4a). Experiments were conducted in casamino acids, or glucose-containing medium, as under conditions where pHV3 is essential (e.g., growth on xylose), segregation defects may be masked by slow growth. In glucose, where the secondary chromosome is less critical, copy number differences between wild-type and *hpaAB* deletion strains were more readily detectable. Deletion of *hpaB* resulted in a significant reduction of the pHV3-to-chromosome ratio, whereas deletion of *hpaA* did not produce a detectable change. This suggests that HpaB functions as the primary driver of pHV3 segregation, directly ensuring proper partitioning and maintenance of the secondary chromosome. In contrast, HpaA may play a modulatory role, contributing to its dynamic localization, but is not essential for maintaining overall pHV3 copy number under the conditions tested. Additionally, subtle or condition-dependent effects of HpaA may not be apparent in glucose-containing medium, where pHV3 is less critical, whereas the role of HpaB is robust and readily observable. Furthermore, we cannot rule out that there could be a redundancy between the HpaA proteins encoded in the *H. volcanii* genome.

Nevertheless, our data show that HpaA is required for the non-random, equidistant distribution of HpaB foci. Without HpaA, HpaB-foci cluster irregularly, potentially reducing fidelity during division. Equipositioning ensures that each daughter cell inherits chromosome copies, particularly critical in pleomorphic disk-shaped cells where the division plane is not fixed by polarity but rather a consequence of the cell shape ^116^. This contrasts with many bacterial ParA-ParB systems, where chromosomes and plasmids are segregated toward specific cellular regions, such as the poles ^113,117^. In *H. volcanii*, equipositioning allows for spatial segregation across two dimensions, accommodating the organism’s polyploidy and pleomorphic morphology. The equidistant pattern may be especially beneficial during late-log phase when cells are disk-shaped and division axes vary.

The dynamic interplay between HpaA and HpaB is conceptually reminiscent of both McdAB and ParAB systems. Like McdA, HpaA may form gradients regulated by local interactions with HpaB foci. The HpaAB system exhibits several features consistent with a Brownian ratchet mechanism. HpaB forms localized foci that are positioned to maximize spacing across the nucleoid, indicating that movement is non-random and spatially regulated. HpaA is dynamic, and its localization likely depends on ATP binding and hydrolysis, as shown by mutations that specifically disrupt these activities. Together, these observations suggest that HpaAB uses diffusion coupled to ATP-driven cycles of binding and release to actively partition the secondary chromosome. This behaviour is reminiscent of the McdAB carboxysome positioning system, in which McdB-bound carboxysomes are similarly distributed across the nucleoid through interactions with the ATPase McdA^38,85,114^. Both systems appear to employ a comparable strategy of equipositioning via localized binding and dynamic ATPase activity. Deletion of *hpaA* results in a dramatic increase in the number of HpaB foci, from approximately average 2–4 in wild type cells to an average of 8-9, and a maximum of 15–20 in *ΔhpaA* strains, roughly corresponding to the total number of pHV3 replicons. This suggests that in wild type cells, HpaA is required to corral multiple replicons into shared HpaB foci, whereas in its absence, each replicon forms an independent HpaB focus. These observations suggest that HpaA acts as both a coraller, promoting the clustering of HpaB-bound replicons, and a spatial organizer, biasing their distribution over the nucleoid. Thus, the behaviour of HpaAB is conceptually highly similar to the McdAB system, where McdA ATPase activity organizes McdB-bound carboxysomes into evenly spaced clusters, highlighting a conserved strategy for the spatial organization of large cellular components via ATP-dependent dynamic interactions.

### Functional implications and evolutionary context

The HpaAB system in *H. volcanii* provides an intriguing example of a hybrid segregation mechanism involving a bacterial origin ParA-like ATPase and an archaeal-specific DNA adaptor, similar to the SegAB system from *S. solfataricus* ^41,42^. This modular system likely evolved via horizontal gene transfer and adaptation to hypersaline conditions, where protein surface charge and LLPS properties are crucial ^118,119^.

The fact that the system enables an equidistant distribution of chromosomes throughout the cytoplasm is likely critical for chromosome inheritance in a polyploid archaeon that is not known to rely on traditional segregation mechanisms. This model of equidistant, repulsion-driven plasmid organization may reflect a broader evolutionary strategy in pleomorphic Archaea to cope with their unique cellular geometry and extreme environments. The fragmented genome of *H. volcanii* requires several segregation systems that ensure stable segregation of the individual replicons. We found that all secondary chromosomes (pHV1-3) contain one HpaAB pair, while the pHV4 replicon contains two of these systems. Since pHV4 is integrated into the main chromosome in the lab strain H26, we think it might be possible that these systems may contribute to the segregation of the main chromosome in *H. volcanii*.

In summary, the HpaAB system shows mechanistic and evolutionary divergence from canonical plasmid partitioning systems, combining elements of both spatial regulation (via HpaA gradients) and phase-separated anchoring (via HpaB foci). It represents the first active chromosome segregation system identified in a highly polyploid archaeon, underscoring the versatility of segregation mechanisms across life domains. Further biochemical and phylogenetic analyses are needed to dissect the ATP cycle of HpaA and its interaction with HpaB, and to trace the evolutionary origin of this modular system.

Recent studies on ParA family ATPases show that DNA-binding is mediated by a conserved array of positively charged residues on the dimer surface, independent of specific DNA motifs ^29^. Mutation of these interaction surfaces disrupts gradient formation and relieves cargo positioning, as observed in MipZ and ParA proteins ^81^. Byrne et al. (2025)^110^ further demonstrated that weakened binding of the adaptor and the ATPase-partner shifts the system from structured equipositioning to mere localization without partitioning; which we also see in the case of HpaB where truncation of the N-terminus leads to disordered aggregation of foci. Complementarily, work in *Magnetospirillum* showed that correct protein gradients, not fixed docking at poles, govern spatial organization ^112^. These insights support our hypothesis that in *H. volcanii*, HpaA forms a dynamic cytosolic gradient that repels HpaB–plasmid complexes into an evenly spaced arrangement, thereby achieving spatial segregation in two dimensions, rather than directed partitioning to poles.

## Acknowledgements

The authors thank all members of the Bramkamp lab for critical discussions. We acknowledge funding from the DFG (BR2915/6-2) and by the International Max Planck Research School for Evolutionary Biology (IMPRS EvolBio) (fellowship to K.P).

