## Supplemental Material Text and Figures for "Equipositioning of Chromosomes in the Polyploid Archaeon *Haloferax volcanii* by HpaAB"

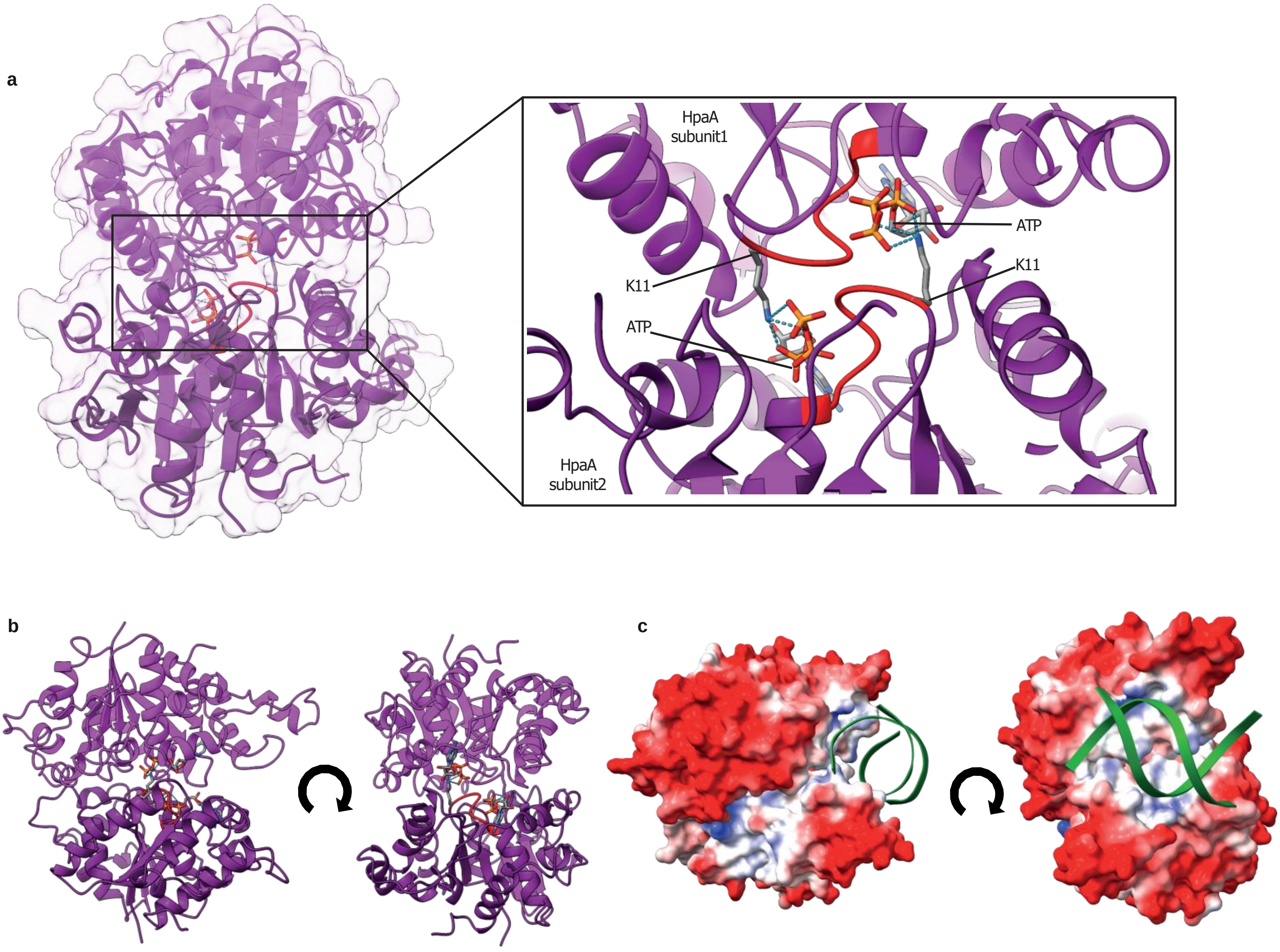


**Supplementary figure 1: HpaA predicted dimer is an ATPase motif sandwich.**
(a) Predicted HpaA dimer structure using AlphaFold 3 showing the conserved Walker A motif in red and ATP inside a sandwich, with zoomed in view at the interaction of the signature lysine K11 from each subunit with ATP of the other subunit. (b and c) Predicted structure (rotated 90°) of the dimer showing majorly negatively charged surface (red), and a positive region where the DNA (green) might bind (blue).

a

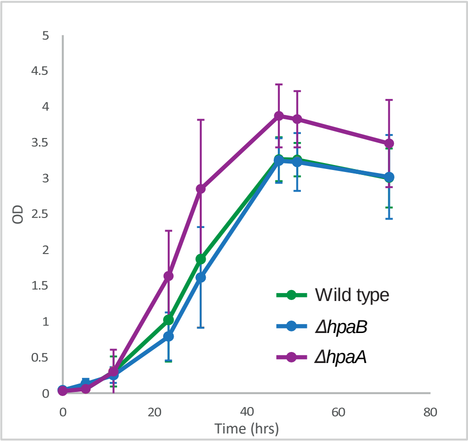


**Supplementary figure 2: Deletion of *hpaA* or *hpaB* does not affect cell viability on glucose.**
(a) Cells were grown under optimal conditions at 42° C, 160 RPM, and 2.1 M NaCl containing synthetic media supplemented with 20 mM glucose. The growth curves show a high degree of variation in growth among different replicates, but all strains remain viable with no severe growth defects.

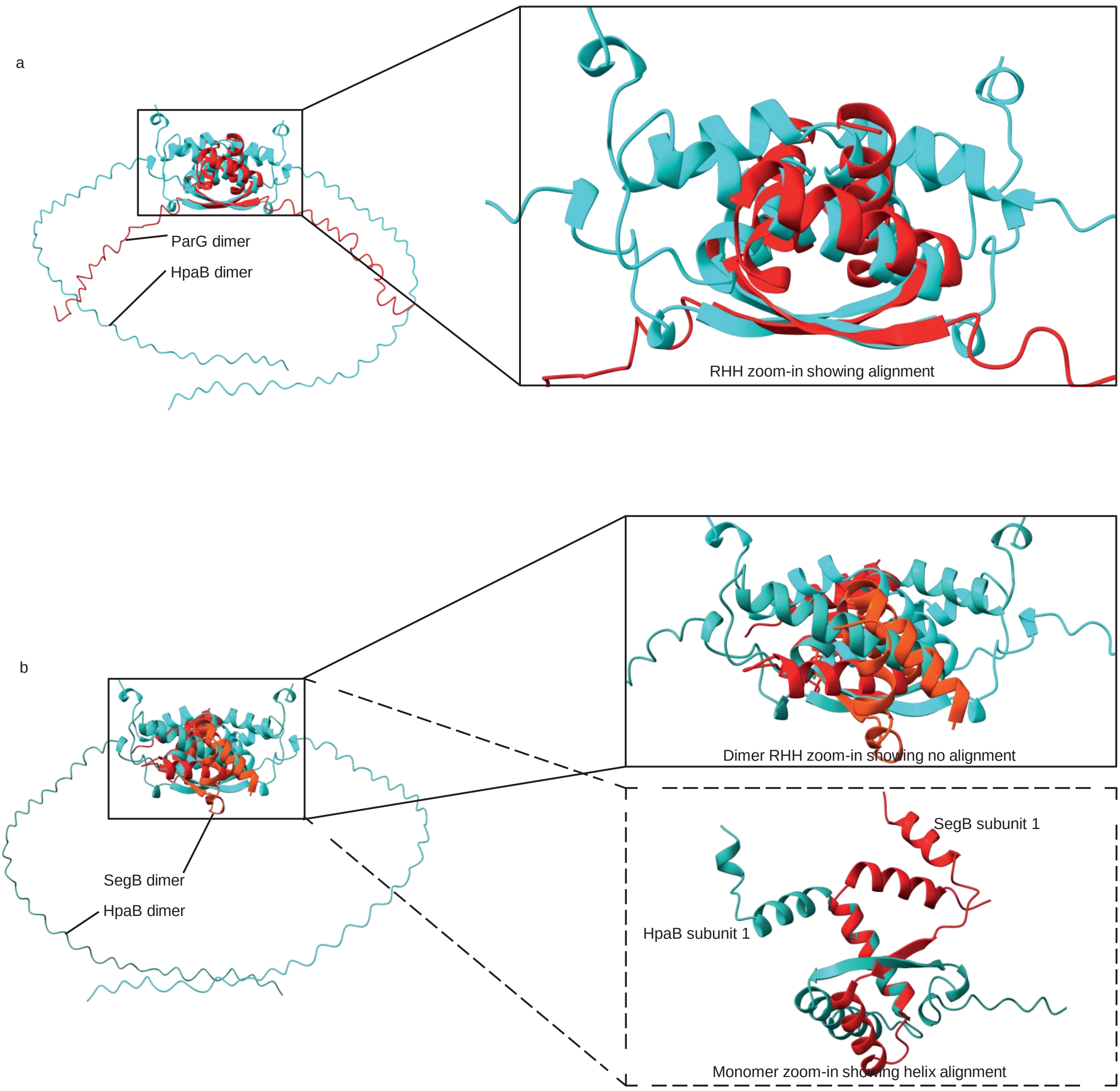


**Supplementary figure 3: HpaB contains a conserved RHH fold**

Comparison of Alphafold 3 predicted structure of HpaB in cyan with the known structures of ParG (PDB: 1P94)[1] and SegB (PDB: 7DUV)[2] in red. (A) Comparison with ParG with inset zoomed-in RHH motif shows similar RHH fold overlap. (B) Comparison with SegB with inset zoomed-in dimer and monomer showing overlapping helix from the RHH motif.

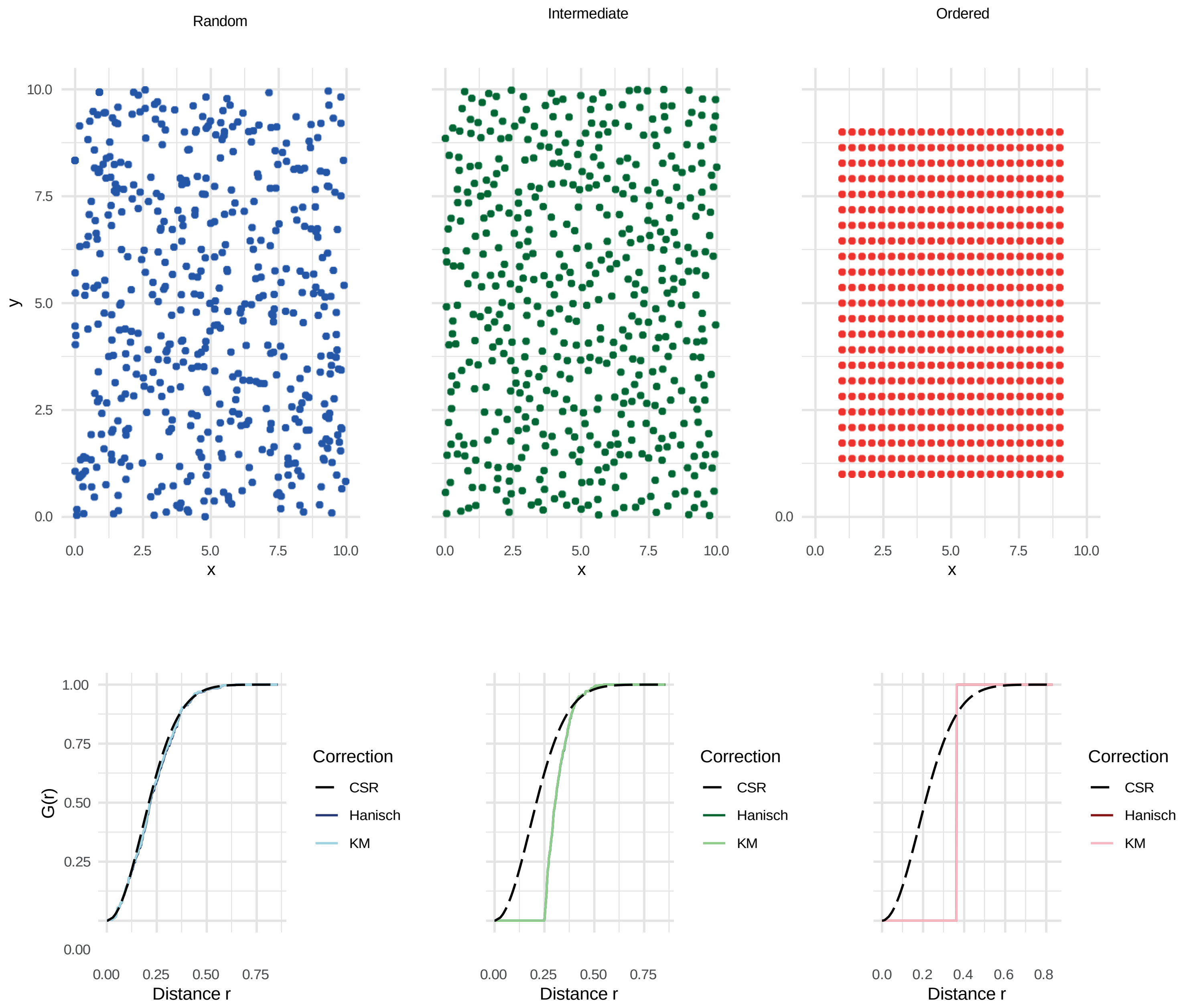


**Supplementary figure 4: Simulated representation of point distribution and G function curves**

The points on a plane in a random, intermediate, or ordered distribution showing clusters and repelling behaviours in the same plane for random (blue), decreased randomness in intermediate (green), and equipositioning in ordered (red). The G-function curve with border corrected nearest neighbour cumulative curves using Hanisch[3] (bold) and km[4] estimates (not-bold). The non-continuous black curve indicates complete spatial randomness on the given set of data; the corrected curves show how the G-function would look for random distribution (overlapping estimates with CSR), intermediate distribution (compression of estimates to a repelling behaviour), and ordered distribution (compressed estimates leading to a small range of distance on x- axis).

**Supplementary video 1:**

Representative fluorescence microscopy time lapse video of oscillation of HpaA−msfGFP in *H. volcanii* H26 wild type cells (red hot intensity). Images were taken 20 times every 6 minutes, and pole to pole oscillation of higher intensity clouds is clearly visible in frames 4 to 6, showing a periodicity of 18−24 minutes for HpaA. Scale bar = 2 µm.

**Supplementary video 2:**

Representative fluorescence microscopy time lapse video of HpaA−msfGFP (red hot intensity) in *H. volcanii* H26 *ΔhpaB* cells. Images were taken 20 times every 6 minutes, and while the protein is dynamic, pole to pole oscillation is not visible. Scale bar = 2 µm.

**Supplementary Table 1: Strains used in this study**

| Strain Name | Description | Reference |
|---|---|---|
| | *Haloferax volcanii* | |
| | H26 Δ*pyrE2* | [5] |
| HPHfx007 | H26 Δ*pyrE2* Δ*hpaB* (pop−out) | This study |
| HPHfx010 | H26 Δ*pyrE2* Δ*hpaB* pTA963 HpaB | This study |
| HPHfx011 | H26 Δ*pyrE2* Δ*hpaB* pTA963 HpaB−msfGFP | This study |
| HPHfx014 | H26 Δ*pyrE2* pTA963 HpaB−msfGFP | This study |
| HPHfx017 | H26 Δ*pyrE2* Δ*hpaB* pTA963 HpaA−msfGFP | This study |
| HPHfx019 | H26 Δ*pyrE2* pTA963 HpaA−msfGFP | This study |
| HPHfx028 | H26 Δ*pyrE2* Δ*hpaA* (pop−out) | This study |
| HPHfx031 | H26 Δ*pyrE2* Δ*hpaA* pTA963 HpaB−msfGFP | This study |
| HPHfx032 | H26 Δ*pyrE2* Δ*hpaA* pTA963 HpaA−msfGFP | This study |
| HPHfx106 | H26 Δ*pyrE2* Δ*hpaB* pTA963 (empty vector) | This study |
| HPHfx107 | H26 Δ*pyrE2* Δ*hpaA* pTA963 (empty vector) | This study |
| HPHfx108 | H26 Δ*pyrE2* Δ*hpaB* pTA963 HpaA | This study |
| HKP003 | H26 Δ*pyrE2* pHVID HpaA−mTurqoise2 HpaB−mScarlet−I | This Study |
| HKP005 | H26 Δ*pyrE2* pTA963 HpaA−Halo | This study |
| HKP007 | H26 Δ*pyrE2* Δ*hpaB* pTA963 HpaA−Halo | This study |
| HKP010 | H26 Δ*pyrE2* pTA963 Δ−msfGFP H70A | This study |
| HKP015 | H26 Δ*pyrE2* pTA963 HpaB−msfGFP Y74A | This study |
| HKP016 | H26 Δ*pyrE2* pTA963 HpaB−msfGFP H70A Y74A | This study |
| HKP025 | H26 Δ*pyrE2* Δ*hpaB* pTA963 HpaB−msfGFP H70A Y74A | This study |
| HKP026 | H26 Δ*pyrE2* Δ*hpaB* pTA963 HpaB−msfGFP Y74A | This study |
| HKP030 | H26 Δ*pyrE2* pTA963 HpaB−msfGFP ΔN−term (−A12) | This study |
| HKP031 | H26 Δ*pyrE2* Δ*hpaB* pTA963 HpaB−msfGFP ΔN−term (−A12) | This study |
| HKP041 | H26 Δ*pyrE2* Δ*hpaA* pTA963 HpaA−msfGFP G12V | This study |

| | | |
|---|---|---|
| HKP042 | H26 Δ*pyrE2* Δ*hpaA* pTA963 HpaA–msfGFP K16A | This study |
| HKP043 | H26 Δ*pyrE2* Δ*hpaA* pTA963 HpaA–msfGFP D40A | This study |

**Supplementary Table 2: Plasmids used in this study**

| Plasmid Name | Description | Reference |
|---|---|---|
| pTA963 | | [6] |
| | pTA963 HpaA | This study |
| | pTA963 HpaB | This study |
| | pTA963 HpaB–msfGFP | This study |
| | pTA963 $HpaB_{H70A}$–msfGFP | This study |
| | pTA963 $HpaB_{Y74A}$–msfGFP | This study |
| | pTA963 $HpaB_{H70A\ Y74A}$–msfGFP | This study |
| | pTA963 $HpaB_{\Delta N\text{–}term}$–msfGFP | This study |
| | pTA963 HpaA–msfGFP | This study |
| | pTA963 HpaA–halotag | This study |
| | pTA963 $HpaA_{G12V}$–msfGFP | This study |
| | pTA963 $HpaA_{K16A}$–msfGFP | This study |
| | pTA963 $HpaA_{D40A}$–msfGFP | This study |
| pTA131 BEMCS | | [7] |
| | pTA 131 BEMCS Δ*hpaB* | This study |
| | pTA 131 BEMCS Δ*hpaA* | This study |
| pHVID 2 | | [8] |
| pHVID 7 | | |
| pHVID 9 | | |
| | pHVID HpaB–mScarlet–I HpaA–mTurqoise2 | This study |

**Supplementary Table 3: Oligonucleotides used in this study**

| Description | Sequence |
|---|---|
| HVO_B0017frg1_s | CGC TTC TCG GGG ACG ACT ACG |
| HVO_B0017frg1_as | GGA TTA TCC GAA GGT CGC CGA AGG GGT CTT CGC |
| HVO_B0017frg2_s | TCG GCG ACC TTC GGA TAA TCC ACC TCC GTC G |
| HVO_B0017frg2_as | GTA CGC GAC CAT GAG GAG TTC G |
| HVO_B0017s_BspH1 | TGG CCT CAT GAG CGA AGA CCC C |
| HVO_B0017as_BamH1 | GTC CGG GGA TCC GAG TTT CGA CGG |
| HVO_B0017as_+3linker | GGA CCC TCC GCC ACC GCT ACC CCC GCC ACC TTC GGA ATC TTC GTC GTC GAC |
| HVO_A0002as_+3linker | GGA CCC TCC GCC ACC GCT ACC CCC GCC ACC CTC GTT GCG ACG TTC GAG GAC |
| HVO_A0459as_+3linker | GGA CCC TCC GCC ACC GCT ACC CCC GCC ACC GTC GAA CGT CAT CCC GTA GCC |
| msfGFP_s+5linker | GGT GGC GGG GGT AGC GGT GGC GGA GGG TCC ATG GGT ACC CTG CAG ATG AG |
| msfGFP_as+3BamHI | CCA CGA GGA TCC CTA TTT GTA GAG CTC ATC CAT GC |
| T3 | AAT TAA CCC TCA CTA AAG GG |
| T7 | TAA TAC GAC TCA CTA TAG GG |
| msfGFP_NcoI_s | TCG TGG CCA TGG GTA CCC TGC AGA TGA G |
| HVO_B0018frg1_s | GAG GAG GAA GTA CAC GAG CGA GG |
| HVO_B0018frg1_as | GAC GAT TTG GAT GGG GAT ATC CGT ACT ATC G |
| HVO_B0018frg2_s | GAT ATC CCC ATC CAA ATC GTC GAA GAA GGC GAG G |
| HVO_B0018frg2_as | GAC TTC TGT CCG GCG CTC CGA G |
| HVO_B0018s_Pci1 | TCC CCA ACA TGT CAC GTG CAG TC |
| GSG–HaloTag_fwd | GGT GGC CCG AGG ATC GGG AGA AAT CGG TAC |
| GSG–HaloTag_rev | AGG CCC GCA CTT AGG AAA TCT CCA GAG TAG AC |
| pTA963_fwd | GAT TTC CTA AGT GCG GGC CTC TTC GCT ATT AC |
| pTA963_rev | CTC CCG ATC CTC GGG CCA CCT CGC CTT C |
| HVO_B0018as_+3linker | GGA CCC TCC GCC ACC GCT ACC CCC GCC ACC TCG GGC CAC CTC GCC TTC TTC |
| HpaB in mScarlet fw | ATACATATGAGCGAAGACCCCTTC |
| HpaB in mScarlet rev | TATGGATCCTTCGGAATCTTCGTCGTC |

| | |
|---|---|
| HpaA in mTq fw | ATACATATGTCACGTGCAGTCAGCG |
| HpaA in mTq rev | TATGGATCCTCGGGCCACCTCGCC |
| pTA963−Halo−Fw | GGT GGC GGG GGT AGC |
| pTA963−Halo−Rev | GTG GTG GTG GTG GTG G |
| pHVID_mTq_fwd<br>pHVID_mTq_rev | GGC CCG AGG CGG CGG GGG CTC GGG G<br>CGT GAC ATG GAT CCC ATA TGT CTA GAA GGT CCG CGA ATG TGA AGT A |
| HpaA_fwd | TGG GAT CCA TGT CAC GTG CAG TCA GCG TC |
| HpaA_rev | CCG CCG CCT CGG GCC ACC TCG CCT TC |
| pHVID_mScarletI_fwd | TTC CGA AGG CGG CGG GGG CTC GGG G |
| pHVID_mScarletI_rev | CGC TCA TGG ATC CCA TAT GTC TAG AAG GTC CGC GAA TGT GAA GTA |
| HpaB_fwd | TGG GAT CCA TGA GCG AAG ACC CCT TC |
| HpaB_rev | CCG CCG CCT TCG GAA TCT TCG TCG TC |
| HpaB H70A OE fw | GAGACCGTCGCCAAGGCGCTG |
| HpaB H70A OE rv | GCGCCTTGGCGACGGTCTCGTC |
| HpaB Y74A OE fw | GGCGCTGGCCGTGCTCGAAG |
| HpaB Y74A OE rv | CTTCGAGCACGGCCAGCGCCTTG |
| pTA963_OLEG_fw | AGATTCCGAAGGTGGCGGGGGTAGCGGT |
| pTA963_OLEG_rv | CTTCGCTCATGTGGTGGTGGTGGTGGTGC |
| HpaB_OLEG_fw | CCACCACCACATGAGCGAAGACCCCTTC |
| HpaB_OLEG_rv | CCCCGCCACCTTCGGAATCTTCGTCGTC |
| HpaB_ΔNTerm_Fw | ATG GGA AGC GGC GAG |

### Supplementary table 4: Statistics Values

| **qPCR comparisons between wild type, Δ*hpaA,* Δ*hpaB,* and complements – Figure 4** | | |
|---|---|---|
| **Comparison** | **p-value (significance) [a]** | **VDA.m (eff. size) [b]** |
| H26/ Δ*hpaB* | 5.74e−14 (****) | 1.000 (large) |
| H26/ Δ*hpaA* | 0.4254(ns) | - |
| Δ*hpaB*_c/ Δ*hpaB* | 0.0017(**) | 0.9950 (large) |
| Δ*hpaB*_c/ H26 | 0.0049(**) | 0.9676 (large) |
| Δ*hpaA*_c/ Δ*hpaA* | 0.0189 (*) | 0.8440 (large) |
| Δ*hpaA*_c/ H26 | 0.1841 (ns) | - |

[a] Adjusted p value and associated significance levels: non-significant (ns), <0.05 (*), <0.01 (**), <0.001 (***), <0.0001(****)

[b] Vargha and Delaney's A and associated effect size: VDA<0.56 (-), 0.56≤VDA<0.64 (small), 0.64≤VDA<0.71 (moderate), 0.64≤VDA (large)
